# Defying Gravity: *WEEP* promotes negative gravitropism in *Prunus persica* (peach) shoots and roots by establishing asymmetric auxin gradients

**DOI:** 10.1101/2023.05.26.542472

**Authors:** Andrea R. Kohler, Andrew Scheil, Joseph L. Hill, Jeffrey R. Allen, Jameel M. Al-Haddad, Charity Z. Goeckeritz, Lucia C. Strader, Frank W. Telewski, Courtney A. Hollender

## Abstract

Trees with weeping shoot architectures are valued for their beauty and serve as tremendous resources for understanding how plants regulate posture control. The *Prunus persica* (peach) weeping phenotype, which has elliptical downward arching branches, is caused by a homozygous mutation in the *WEEP* gene. Until now, little was known about the function of WEEP protein despite its high conservation throughout Plantae. Here, we present the results of anatomical, biochemical, biomechanical, physiological, and molecular experiments that provide insight into WEEP function. Our data suggest that weeping peach does not have defects in branch structure. Rather, transcriptomes from the adaxial (upper) and abaxial (lower) sides of standard and weeping branch shoot tips revealed flipped expression patterns for genes associated with early auxin response, tissue patterning, cell elongation, and tension wood development.

This suggests that WEEP promotes polar auxin transport toward the lower side during shoot gravitropic response, leading to cell elongation and tension wood development. In addition, weeping peach trees exhibited steeper root systems and faster root gravitropic response, just as barley and wheat with mutations in their *WEEP* homolog *EGT2*. This suggests that the role of WEEP in regulating lateral organ angles and orientations during gravitropism may be conserved. Additionally, size-exclusion chromatography indicated that WEEP proteins self-oligomerize, like other SAM-domain proteins. This oligomerization may be required for WEEP to function in formation of protein complexes during auxin transport. Collectively, our results from weeping peach provide new insight into polar auxin transport mechanisms associated with gravitropism and lateral shoot and root orientation.

## Introduction

Weeping trees have long been prized for their aesthetic beauty and unique shape. This pendulous growth habit, where branches bend or grow downward in the direction of gravity, exists in both gymnosperm and angiosperm lineages. The weeping trait has been mapped to single, but distinct, loci in multiple species, including Eastern redbud (*Cercis canadensis*), morning glory (*Pharbitis nil*), Japanese apricot (*Prunus mume*), and peach (*Prunus persica*) (Kitazawa et al., 2005; Hollender et al., 2018; Chen and Werner, 2021; Li et al., 2021b). Despite often being controlled by a single locus, the change to a pendulous growth habit leads to diverse alterations in plant anatomy and physiology—such as modifications in light interception, canopy density, and canopy size. Studying the genes which control weeping traits can provide insights into the molecular mechanisms by which branch orientation is regulated, many of which are still unknown. This will ultimately benefit production strategies for diverse crop species, as control of branch orientation is crucial to aspects such as planting density, spray coverage, yield (Ku et al., 2011; Zhao et al., 2014; Roychoudhry and Kepinski, 2015; Xu et al., 2017; González-Arcos et al., 2019).

Here, we investigate the function of *PpeWEEP,* the causative gene for a weeping peach architecture. The branches of peach trees with a homozygous *WEEP* deletion grow downwards in an elliptical trajectory beginning early in development (Figure 1A-D). This phenotype is visible within the first phytomer of primary shoots and branches (Figure 1E and F) and continues throughout their life cycle (Hollender et al., 2018). In addition, weeping peach shoots do not exhibit negative gravitropic responses. Their shoots do not reorient upward after being rotated 90° or 180° (upside-down), and growth from existing and new shoots following reorientation arches downwards (Figure 1G-I; Hollender et al., 2018).

**Figure 1:**
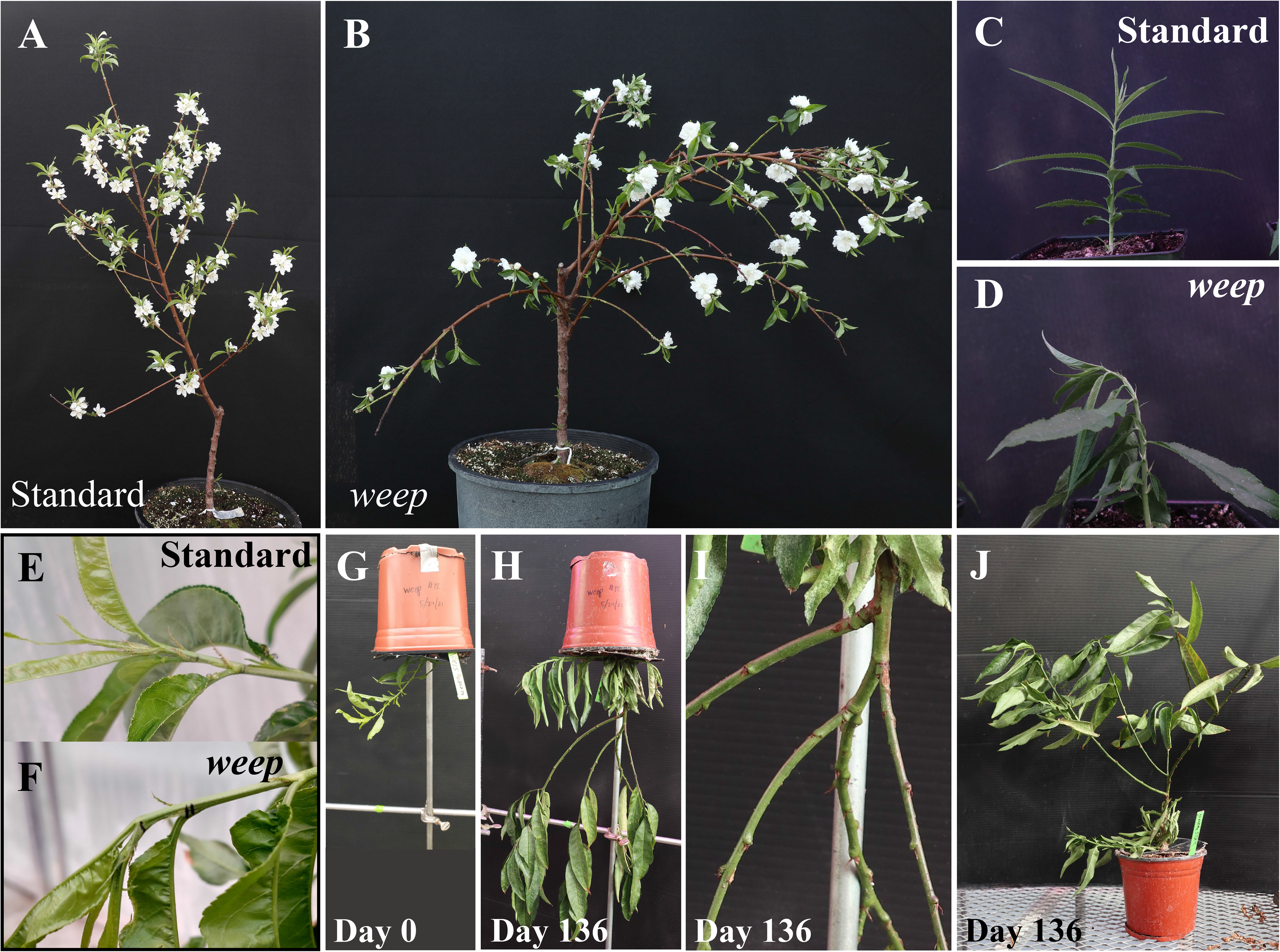
Standard and weeping peach phenotypes. Adult tree phenotypes (A-B), seedling phenotypes (C-D), shoot tip phenotypes in adult trees (E-F) weeping tree reoriented 180° (G), the same tree at 136 days after rotation (H-I), weeping tree restored to upright orientation on day 136 (J).

The WEEP protein sequence is highly conserved throughout vascular plant clades, suggesting it plays an essential role in plant development (Hollender et al., 2018). *WEEP* codes for a small protein (125 amino acids) of unknown function that has a sterile alpha motif (SAM) domain which, at 68 amino acids, constitutes over half of the protein (Hollender et al., 2018). SAM domains are versatile interaction domains found throughout eukaryotes that can bind proteins, RNA, or lipids (Qiao and Bowie, 2005; Denay et al., 2017). SAM domains frequently function in the formation of protein homo- or hetero-oligomers or polymers (Qiao and Bowie, 2005; Denay et al., 2017). Polymerization occurs head-to-tail through association of two conserved interaction regions, the negatively charged mid-loop and the positively charged end helix (Sayou et al., 2016; Denay et al., 2017). The formation of oligomers or polymers is often essential for protein function via increasing stability, altering binding strength, or otherwise regulating protein function (Qiao and Bowie, 2005; Denay et al., 2017). In *Arabidopsis thaliana* (arabidopsis), 12 genes containing SAM domains have been identified, including the *PpeWEEP* homolog *AtWEEP* (AT3G07760, also known as *SAM5*; Denay et al., 2017; Hollender et al., 2018). SAM domain functions have been characterized in three arabidopsis proteins: LEAFY (LFY), TRNA IMPORT COMPONENT 1 (TRIC1), and TRIC2. TRIC1 and TRIC2 are mitochondrial tRNA importers, whose SAM domains both bind tRNA and enable homopolymerization of TRIC1 and heteropolymerization of TRIC1 and TRIC2 (Murcha et al., 2016; Denay et al., 2017). LFY is a floral identity transcription factor, whose SAM domain mediates self-oligomerization required for it to bind to low-affinity DNA binding sites and closed chromatin (Sayou et al., 2016; Denay et al., 2017). Thus, in all plant proteins where they have been characterized, SAM domains enable protein oligomerization and are essential for protein function.

Due to the broad utility of a binding domain in proteins of highly varied cellular functions, the presence of a SAM domain provides only limited clues to WEEP function in plant physiology and development. However, studies in woody and herbaceous plants suggest *WEEP* has a conserved function in regulating lateral organ orientations. Plum (*Prunus domestica*) trees with reduced *WEEP* expression have branches that wander and arch (Hollender et al., 2018). In contrast, *weep* loss of function mutants in arabidopsis, wheat (*Triticum aestivum*), and barley (*Hordeum vulgeare*) have normal shoot architectures but steeper root systems, due to narrower seminal and lateral root angles (Hollender et al., 2018; Kirschner et al., 2021; Johnson et al., 2022). Mutations in the barley and wheat genes, named *ENHANCED GRAVITROPISM* 2 (*EGT2*), also cause accelerated root gravitropism responses (Kirschner et al., 2021; Guo et al., 2023).

Other work has suggested *WEEP* plays a role in regulating cell expansion. In melon (*Cucumis melo*), the *WEEP* homolog *DOWNWARD LEAF CURLING* (*CmDLC*) was originally identified as being upregulated during fruit expansion (Kee et al., 2009). Overexpression of *CmDLC* in arabidopsis leads to semi-dwarfism and reductions in leaf pavement cell size and number, especially on the abaxial side of the leaf (Kee et al., 2009). In arabidopsis, *AtWEEP* is upregulated in developing leaves when mature leaves are shaded, a treatment which slows the growth of developing leaves (Coupe et al., 2006). In barley roots, mutation in the *WEEP* homolog *EGT2* leads to decreased expression of *EXPANSIN* genes in the root elongation zone (Kirschner et al., 2021; Guo et al., 2023).

Transcriptomics databases show that *WEEP* is expressed throughout plant organs, but is specifically upregulated in tissues consistent with a role in gravitropism, abaxial/adaxial polarity, and lateral organ development. *AtWEEP* is expressed in the hypocotyl, root, mature leaves, flowers, and seeds (Klepikova et al., 2016). Despite this ubiquitous expression, *AtWEEP* is differentially regulated in different tissues. In the shoot apex, it is upregulated in the enlarged peripheral zone of the meristem, in the organ boundary, on the adaxial side of leaf primordia, and in the epidermis (Tian et al., 2019). This localization may be related to gibberellin signaling and tissue patterning, as *AtWEEP* is transcriptionally regulated by the DELLA protein GAI, in the presence of CUP-SHAPED COTYLEDON2 (CUC2), which is essential for organ boundary specification (Barro-Trastoy et al., 2022). In the root, *AtWEEP* is highly upregulated in the endodermis, the columella, and the stele (Ryu et al., 2019).

Thus far, protein interaction candidates identified for WEEP homologs are involved in cell wall synthesis or membrane transport. In transient heterologous expression studies, WEEP protein homologs have localized to the plasma membrane (CmDLC in onion) or the nucleus and cytoplasm (barley EGT2 in tobacco; Kee et al., 2009; Guo et al., 2023). In barley, three candidate protein interactors for EGT2 were identified with yeast-two-hybrid screening and confirmed with Bimolecular Fluorescence Complementation – GXM, OMT, and HMT (Guo et al., 2023). GXM is a glucuronoxylan methyltransferase, with sequence similarity to the three arabidopsis GXMs (GXM1,2, and 3; Guo et al., 2023). In arabidopsis, these are responsible for methylation of glucuronic acid residues in xylan (Yuan et al., 2014). OMT is a homolog of AtOMT1, an oxygen methyl-transferase that methylates both 5-hydroxyconiferaldehyde and 3,4- dihydroxyphenyl compounds during the production of syringyl (S) lignin (Nakatsubo et al., 2008; Guo et al., 2023). Interestingly, increased syringyl to guaiacyl lignin ratio has observed to cause more rapid gravitropic response in poplar (Al-Haddad et al., 2009). HMT shows homology to proteins in the heavy metal transport/detoxification superfamily (Guo et al., 2023). In addition, in arabidopsis, the AtWEEP protein was identified to interact with CPK13, a nuclear- and plasma membrane-localized kinase which acts in a calcium-independent manner to inhibit the voltage-dependent K^+^ channel KAT2, which is important for light-induced stomatal opening (Jones et al., 2014; Ronzier et al., 2014; Simeunovic et al., 2016; Denay et al., 2017). CPK13 also phosphorylates the heat-shock factor AtHSfB2a (Simeunovic et al., 2016).

The pendulous habit of *weep* peaches may be due to reduced structural integrity, an impaired ability to sense or respond to gravity, or a positive gravitropic response. Lack of structural integrity leads to downward bending through self-loading, as the branch is unable to support its own weight. Supporting the structural integrity hypothesis, reduced xylem tissue width and delayed development of tension wood are associated with a weeping architecture in Japanese cherry (*Prunus spachiana*; Nakamura *et al*., 1994; Baba *et al*., 1995; Yoshida *et al*., 1999). Tension wood refers to localized changes in secondary cell wall composition which occur within the upper side of angiosperm shoots in response to gravistimulation or biomechanical stress. It is associated with an increase in the proportion of crystalline cellulose, decreases in microfibril angle, and sometimes an increase in S:G ratio of lignin monomers or the formation of gelatinous fibers (Qiu et al., 2008; Felten and Sundberg, 2013). Tension wood generates internal forces that return branches back towards their original position or “set-point angle” (Felten and Sundberg, 2013; Roychoudhry et al., 2013). Application of GA to the apical bud of *P. spachiana* rescued this weeping phenotype through increase in xylem width and earlier formation of tension wood (Nakamura et al., 1994; Taniguchi et al., 2017). This effect was not observed in *weep* peaches, where GA application did not alter the weeping phenotype (Hollender et al., 2018). Similar to Japanese cherry, weeping Japanese apricot (*Prunus mume*) trees have decreased xylem tissue width, and lack phloem fibers (Li et al., 2021b).

Alternatively, the weeping peach phenotype could be due to impaired shoot gravitropism, indicating a role for *WEEP* in the gravitropic signaling pathway. This pathway can be divided into five main stages. First, the gravity vector is sensed by statocytes (endodermal cells in shoots and columella cells in roots), which contain amyloplasts that serve as statoliths. Second, polarity is established through vesicle trafficking and PIN-FORMED (PIN) auxin efflux carrier localization (Nakamura et al., 2019; Zhang et al., 2020). Third, asymmetric auxin transport establishes an auxin gradient with higher auxin levels on the lower side of the shoot or root (the Cholodny-Went hypothesis; Moore, 2002; Rakusová et al., 2011). Fourth, localized expression of rapid auxin response genes such as *SMALL AUXIN UPREGULATED RNAs* (*SAURs*) leads to asymmetric growth and bending through acidification of the apoplast, leading to cell division and elongation (acid growth hypothesis; Spartz et al., 2012; Ren and Gray, 2015; Wang et al., 2020; Li et al., 2021a). Defects in any of these stages impair gravitropism, and mutants frequently exhibit weeping shoot phenotypes. Weeping shoot phenotypes have been described in mutants which affect endodermis development (*shortroot*, *scarecrow),* amyloplast sedimentation (*shoot gravitropism, zigzag*), and starchless mutants (*phosphoglucomutase deficient*; Fukaki et al., 1996; Kato et al., 2002; Kitazawa et al., 2005; Morita et al., 2007). Weeping shoot phenotypes have also been observed in arabidopsis *lazy1,2,4* mutants, which do not have correct PIN protein polarization and lack an auxin gradient (Taniguchi et al., 2017; Yoshihara and Spalding, 2017). Thus, agravitropism from defects at any of these points in the gravitropism signaling pathway can cause a weeping phenotype.

Finally, the weeping peach phenotype might result from positively gravitropic shoot growth. Positive shoot gravitropism could occur either through inversion of the gravitropic auxin gradient or inversion of the growth response to auxin. Inversion of the shoot auxin gradient and positive shoot gravitropism (rootward growth) have been observed in the arabidopsis *LAZY1* mutant allele AtLAZY1^1L92A/I94A^, which also exhibits a pronounced weeping phenotype (Yoshihara and Spalding, 2020). Alternatively, the growth response to auxin could be inverted, with auxin inhibiting growth (as in roots) rather than promoting it (as in shoots). The mechanism for auxin inhibition of root elongation is not well understood, and is complicated by the apparently dosage-dependent effects of auxin (Barbez et al., 2017; Du et al., 2020).

Here, we present anatomical, physiological, biomechanical, and molecular characterizations of weeping peach branches to assess the hypotheses for the cause of the weeping phenotype and identify the molecular function of PpeWEEP in branch orientation control. Our results suggest that, in peach, the WEEP protein is not required for structural integrity in branches. Rather, WEEP promotes negative gravitropism in both shoots and roots and plays a crucial role in establishing the gravitropic auxin gradient in shoots.

## Results

### Weeping branches have minimal differences in anatomy and wood composition

To address the hypothesis that the weeping habit in peach was due to deleterious changes in branch structure, our early investigations into *WEEP* function focused on branch anatomy and wood composition. For this, and other comparisons, we utilized clonally propagated individuals from the previously established segregating F2 population of peaches used to identify the *weep* mutation (Hollender et al., 2018). Thus, the *weep* and ‘standard’ plant material come from siblings with similar genetic backgrounds – the main difference being the absence or presence of the *weep* mutation.

Interestingly, when a weeping tree that was grown upside-down for several months was reoriented to vertical, the existing branches remained in position, and did not droop downwards in the direction of gravity, suggesting that the branches are strong enough to maintain an upright position (Figure 1J). To assess branch anatomy, actively growing shoot tips (new growth) and dormant wood from the prior year’s growth were hand-sectioned. Percentages of xylem and pith tissues were calculated relative to the total cross-sectional area. Although it was not always visually obvious, the new growth of weeping branches had a greater percentage of pith and smaller percentage xylem, compared to standard shoot tips at a similar developmental stage (Figure 2, A and B). In contrast, no significant differences between weeping and standard branches were detected in dormant growth, indicating that the differences in xylem width were transient and disappeared by the end of the growing season (Figure 2, A and B).

**Figure 2.**
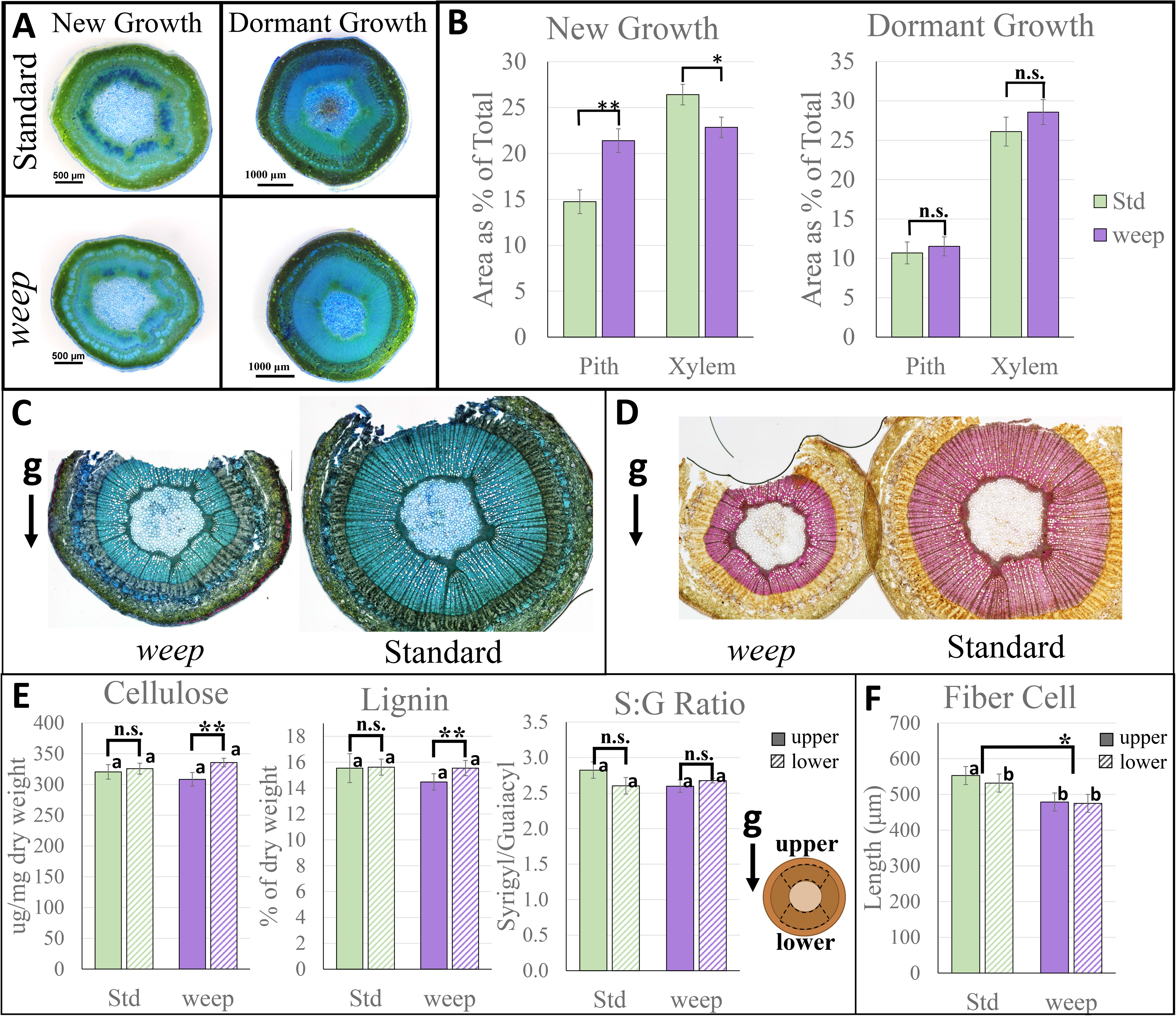
Wood Anatomy and Composition. New growth and dormant first-year growth hand-sectioned and stained with toluidine. (A) Area of pith and xylem relative to total cross-sectional area, brackets show t-tests between genotypes (B). Dormant wood samples stained with toluidine for cellulose (C) and phloroglucinol for lignin (D) to assess tension wood formation. Notches indicate the upper side of the branch. Cell wall polymers in upper and lower standard branches (E). Means with the same letter are not significantly different at α=0.05 in an all pairwise comparison of means with Tukey tests. Bracketed sets indicate the results of paired t-tests between top and bottom of each branch within a genotype. Length of fiber cells from digested wood (F); letters indicate the result of all pairwise comparisons with t-tests, and bracket indicates t-test between genotypes. Std=standard peach, weep=weeping peach. Error bars show standard error. ** indicates significantly different at α=0.05, * indicates significantly different at α=0.10, n.s. indicates not significant at the α=0.10 level.

To investigate wood composition and tension wood formation, lignin and cellulose content were assessed for each genotype, both visually and chemically. Tension wood is characterized by high levels of cellulose, reduced lignin, and often contains obviously visible gelatinous fibers. In contrast, the “opposite wood” on the lower (abaxial) side is enriched in lignin and does not have these specialized fiber cells.

Thin cross sections of dormant one-year old branches were stained with toluidine blue to visualize cellulose content, or phloroglucinol to visualize lignin. No obvious differences in staining intensity or localization were detectible between the two genotypes, nor were gelatinous fibers identifiable (Figure 2, C and D). Biochemical analyses of upper and lower tissues from bisected weeping and standard branches were subsequently performed (Figure 2E; Supplemental Figure S1). No differences in acetyl bromide soluble lignin (ABSL) content, proportion of syringyl (S) and guaiacyl (G) lignin units, or crystalline cellulose content were detected between genotypes or tissue types (Figure 2E). However, the corresponding neutral sugar analysis revealed that weeping peach tissues had lower concentrations of glucose than the standard branch tissues (Supplemental Figure S1).

Lastly, cell size differences between tissues in the upper and lower regions of standard and weeping branches were investigated. Due to the direction of curvature, we hypothesized that epidermal, cortical, and/or xylem cells on the upper side of weeping branches would be longer than those on the lower, and vice-versa in standard branches. No obvious differences in cell size were identified from longitudinal sections, and cell length quantifications proved challenging, partially due to the presence and placement of vegetative lateral buds on branches (Supplemental Figure S2). As an alternative, we addressed cell size by measuring the length of wood fiber cells isolated from macerated tissues from upper and lower portions of bisected standard and weeping peach branches. Fiber cells from the upper tissues of standard peach branches were consistently longer than those from the underside of standard branches (Fig 2F). However, no difference was detected between fiber cell lengths from upper and lower weeping branch tissues. Interestingly, when fiber cell lengths from upper and lower tissues for each genotype were combined, the fiber cells of weeping branches were slightly, but significantly shorter than those from standard branches (p < 0.10; Figure 2F).

### Weeping shoots do not have decreased structural integrity

To determine if weeping peach shoots curve downward because their branches are too weak to support their own weight, we assessed their biomechanical properties. A universal testing machine was used to measure shoot flexibility in the elastic region of its deformation (flexural stiffness-EI), tissue flexibility (modulus of elasticity-MOE), and tissue strength (modulus of rupture—MOR). These tests were performed on actively growing shoot tips because weeping branches already exhibit curvature at this early developmental stage (Figure 1, E and F). Since wood development is a dynamic process, it is impacted by shoot tip orientation as the shoot responds to gravitational forces. Thus, shoots in different orientations may exhibit different mechanical properties. For this experiment, shoots were categorized into “upward”, “downward”, and “outward” orientation (Figure 3A). Only outward shoots were used for statistical comparison between the genotypes because standard trees have only upward and outward shoots, while weeping trees only have outward and downward shoots. No significant differences in EI, MOE, or MOR were observed between standard and *weep* in actively growing, outward-oriented shoot tips (Figure 3B). These biomechanical observations indicate that changes observed in *weep* shoot anatomy do not lead to significant alterations in the shoot structural integrity, and *weep* shoots do not bend in the direction of gravity due to changes in branch stiffness or strength.

**Figure 3.**
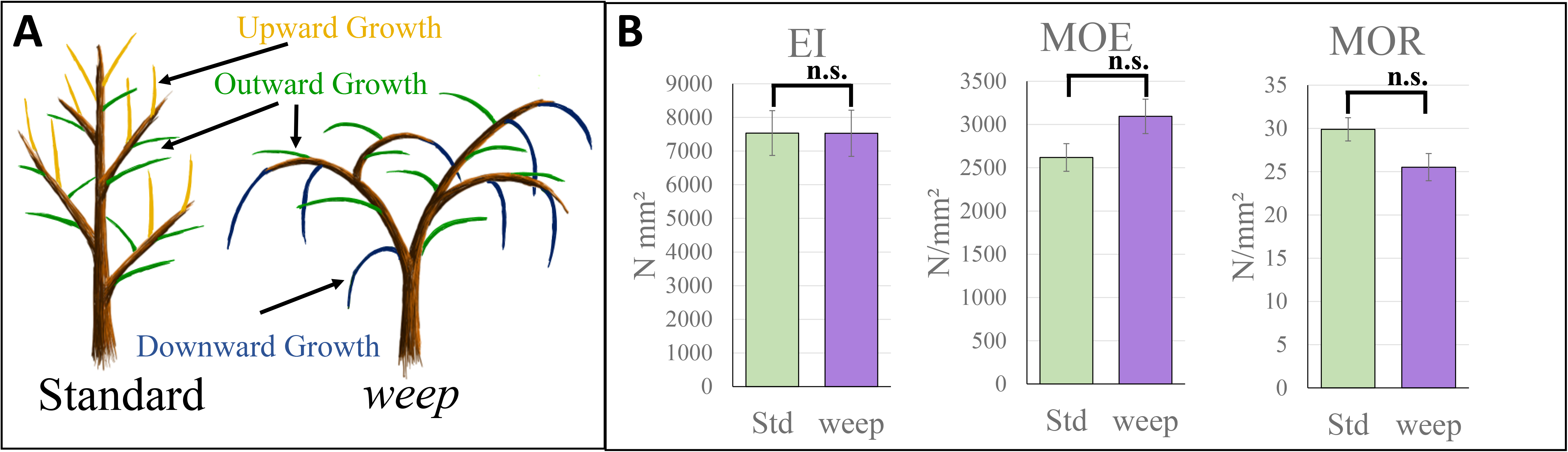
Biomechanical properties of actively growing standard and weeping peach shoots. Actively growing branches of standard (Std) and weeping (weep) peach trees were categorized into upward (gold), outward (green), or downward (blue) orientations (A). Flexural stiffness (EI), the modulus of elasticity (MOE), or the modulus of rupture (MOR) are shown for outward growing shoot tips (B). Bars represent standard error. Pairwise comparisons were done using t-test; n.s. indicates not significant at the 0.1 level.

### Weeping shoot endodermis contains amyloplasts with normal sedimentation

Given the evidence that the weeping phenotype is not due to a loss of structural integrity, coupled with the observation that weeping peach shoots do not bend upwards in response to gravity, we next explored the hypothesis that the *weep* peach mutant has defects in the shoot gravitropism pathway.

To investigate gravitropic perception, we assessed if *weep* mutants had normal amyloplasts and amyloplast sedimentation. Fifty-micron thick fresh vibratome sections of actively growing shoot tips stained with Lugol’s solution revealed that starch-filled amyloplasts were present in both standard and weeping branch shoot endodermis (Figure 4A). Additionally, 1 µm thick sections of resin-embedded shoots tips from outward-oriented branches of each genotype revealed normal amyloplast sedimentation in response to gravity within *weep* shoots. Amyloplasts were consistently found on the lower side of the endodermal cells in both weeping and standard branches (Figure 4B). Thus, weeping shoots contain phenotypically normal amyloplasts and amyloplast sedimentation.

**Figure 4:**
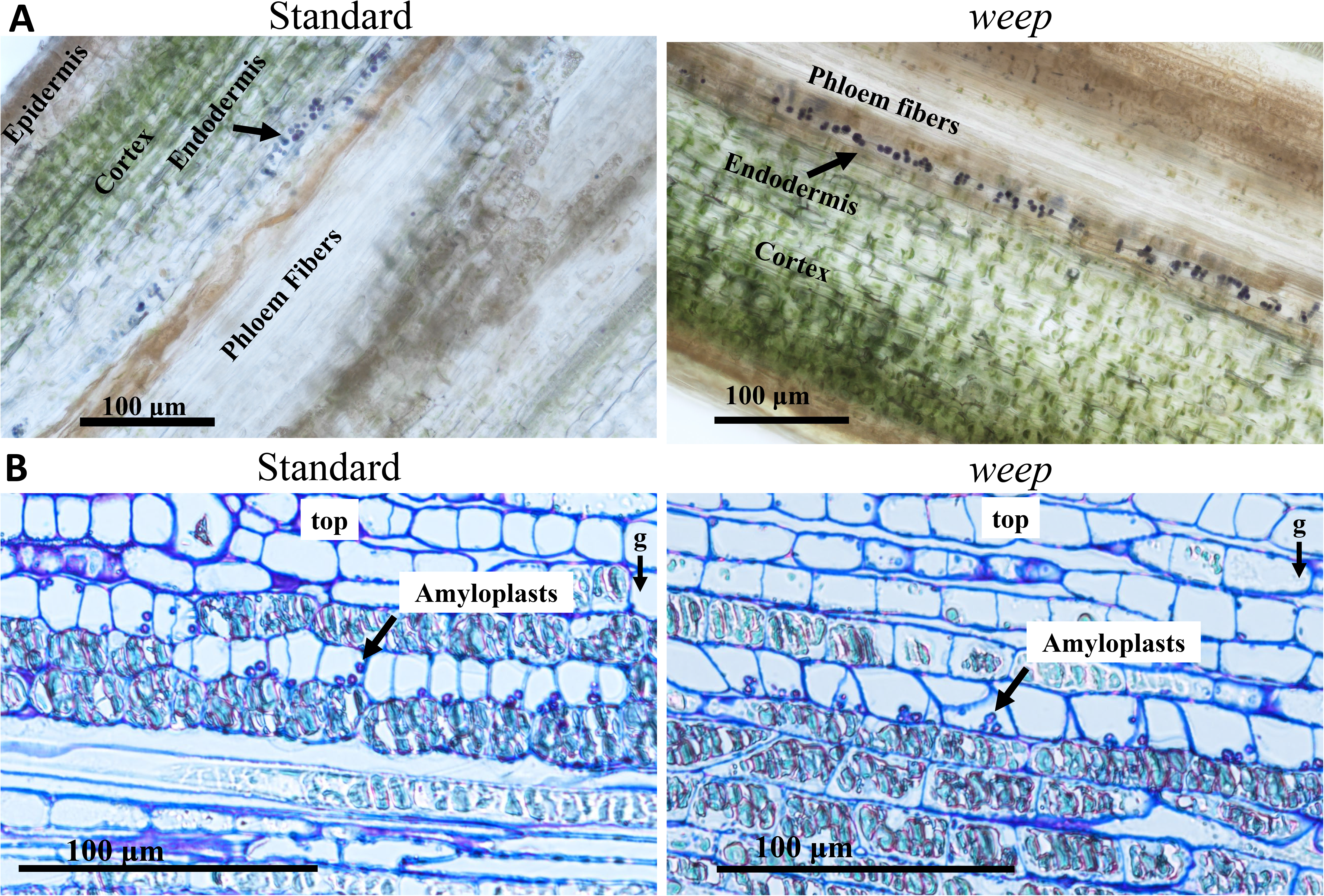
Amyloplast sedimentation. (A) Longitudinal sections of fresh standard and weeping (weep) branches 1cm below the shoot apex stained for starch with Lugol’s solution. (B) Thin sections of resin-embedded standard and weep branches 0.5cm below the shoot apex reveal proper amyloplast sedimentation. Gravity vector indicated by arrow under the g.

### Both standard and weeping shoots bend away from asymmetric exogenous auxin application

We next assessed if weeping shoots were able to exhibit a normal elongation response and bend away from asymmetric auxin localization. Two complementary experiments were performed to simulate the asymmetric auxin concentrations that form as part of the gravitropism response pathway. First, 1% IAA in lanolin paste was applied to the upper or lower side of *weeping* and standard shoots on trees growing in our greenhouse (Figure 5A and B). Second, 1% IAA was applied to one side of upright detached shoots with leaves removed (Figure 5C). In both experiments, weeping and standard shoots bent away from unilateral auxin application, regardless of where it was applied (Figure 5). In both genotypes the degree of bending response was highly variable; however, the direction of bending was consistent across all samples from both genotypes. Therefore, weeping shoots respond normally to auxin, elongating and bending away from areas of high auxin concentration.

**Figure 5:**
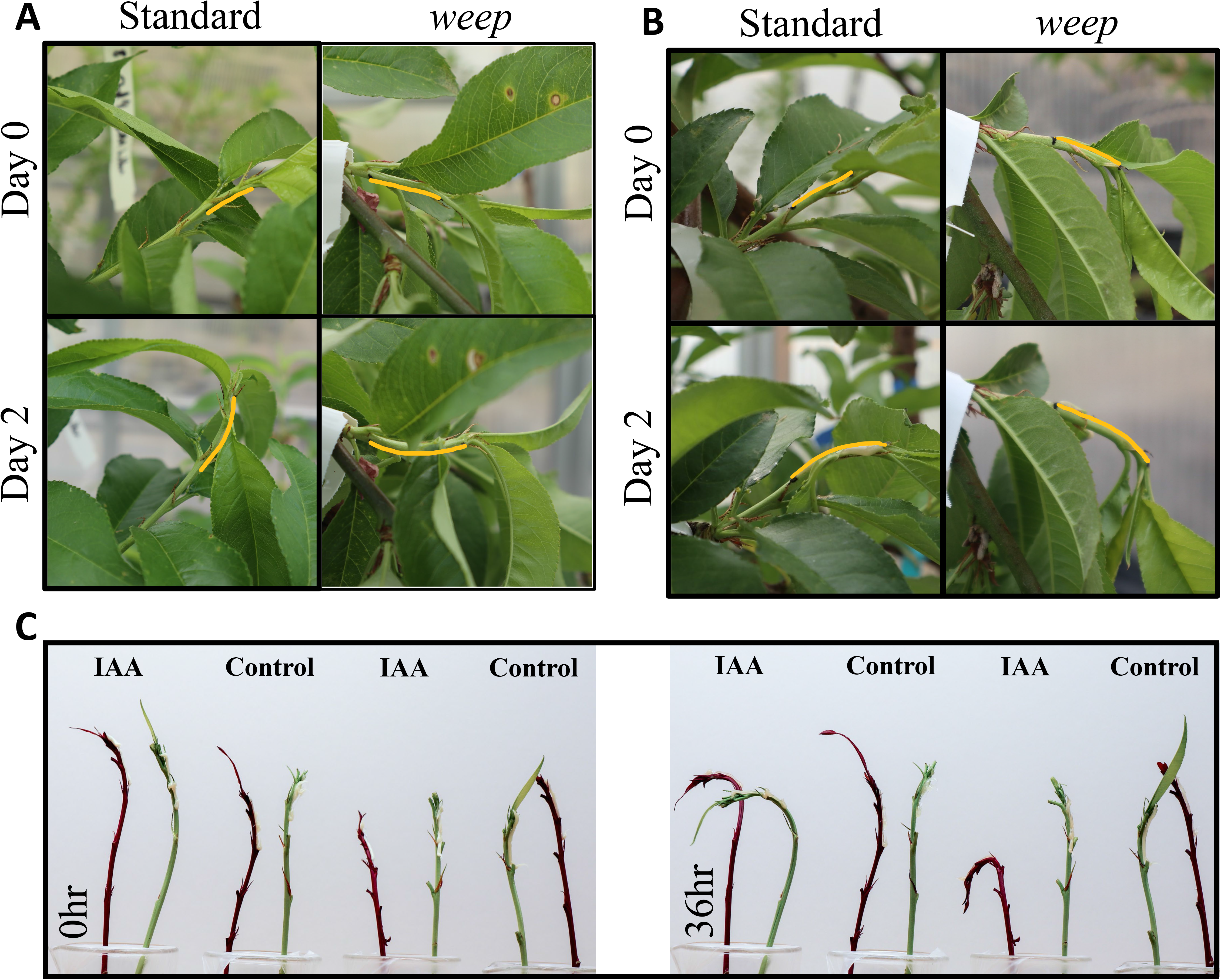
Bending response to unilateral auxin application. Application of 1% IAA in lanolin (yellow line) to the bottom (A) or the top (B) of shoot tips of standard and weeping (weep) trees in the greenhouse. (C) Application of auxin to the right side of upright, detached shoots. Red shoots are from weeping trees that also contain an unlinked anthocyanin phenotype. Green shoots are from standard trees.

### Transcriptome analyses reveal weeping peaches have inverted expression of auxin response and wood development genes

Considering the ability of weeping shoot tips to respond properly to exogenous auxin, we explored the hypothesis that the auxin localization in weeping branches was flipped compared to standard branches. Specifically, we anticipated that standard branch shoot tips would have a greater concentration of auxin in the lower (abaxial) side, and the reverse would be true for weeping peach branches.

To test this, tissues were harvested from the adaxial (upper) and abaxial (lower) sides of the first (IN1) and second (IN2) internodes of actively growing standard and weeping shoot tips (Supplemental Figure S3). Initially, LC/MS/MS was used to measure hormone concentrations of pooled samples consisting of 6-8 shoot tips from the same tree per replicate, for each tissue type. In addition to IAA, concentrations of salicylic acid and ABA were also measured, as they are associated with elongation and tension wood formation, respectively. Surprisingly, no statistically significant differences in IAA concentration were detected between upper and lower tissues for either internode within either genotype (Supplemental Figure S3). Pairwise comparisons between genotypes also showed no significant differences for any of the hormones tested. This may be due to a greater variation in hormone concentration between shoot tips than between the upper and lower tissue within an individual shoot tip. Thus, pooling of shoot tips could have obscured the hormone difference between upper and lower tissues.

To further assess differences between *weep* and standard in polarity of upper and lower tissues, RNA was collected and sequenced. To address problems potentially caused by pooling tissues, RNA was extracted from individual branches. A principal component analysis (PCA) indicated expression profile differences between internodes when all samples were analyzed together (Supplemental Figure S4). Within each internode, the genotypes also formed distinct clusters, particularly in IN2 (Supplemental Figure S5). In addition, differences between upper and lower tissues of a given shoot tip were much smaller than differences between individual shoot tips, confirming our suspicions on why we were unable to detect differences in hormone concentrations (Supplemental Figure S5).

Transcriptional differences were calculated for each genotype between upper and lower tissues from the first two internodes (IN1 and IN2) of outward branch shoot tips (see Sup. Figure S2A). For each internode, genes with a two-fold change in expression and a Bonferroni value < 0.01 in either genotype were selected as genes of interest (GOI; Supplemental Tables S1 and S2). This resulted in 97 GOI for IN1 (Figure 6 and Supplemental Table S1) and 213 GOI for IN2 (Figure 7, Supplemental Figure 6, and Supplemental Table S2). Expression differences between upper and lower branch tissues were more prevalent and stronger in the weeping branches than standard ones (more genes were differentially expressed in the *weep* mutants than in standard, and the fold change for these genes was often larger in *weep*). The greater number of differentially expressed genes (DEGs) in the IN2 samples may be due to adaxial/abaxial polarity being more strongly established further down from the meristem.

**Figure 6:**
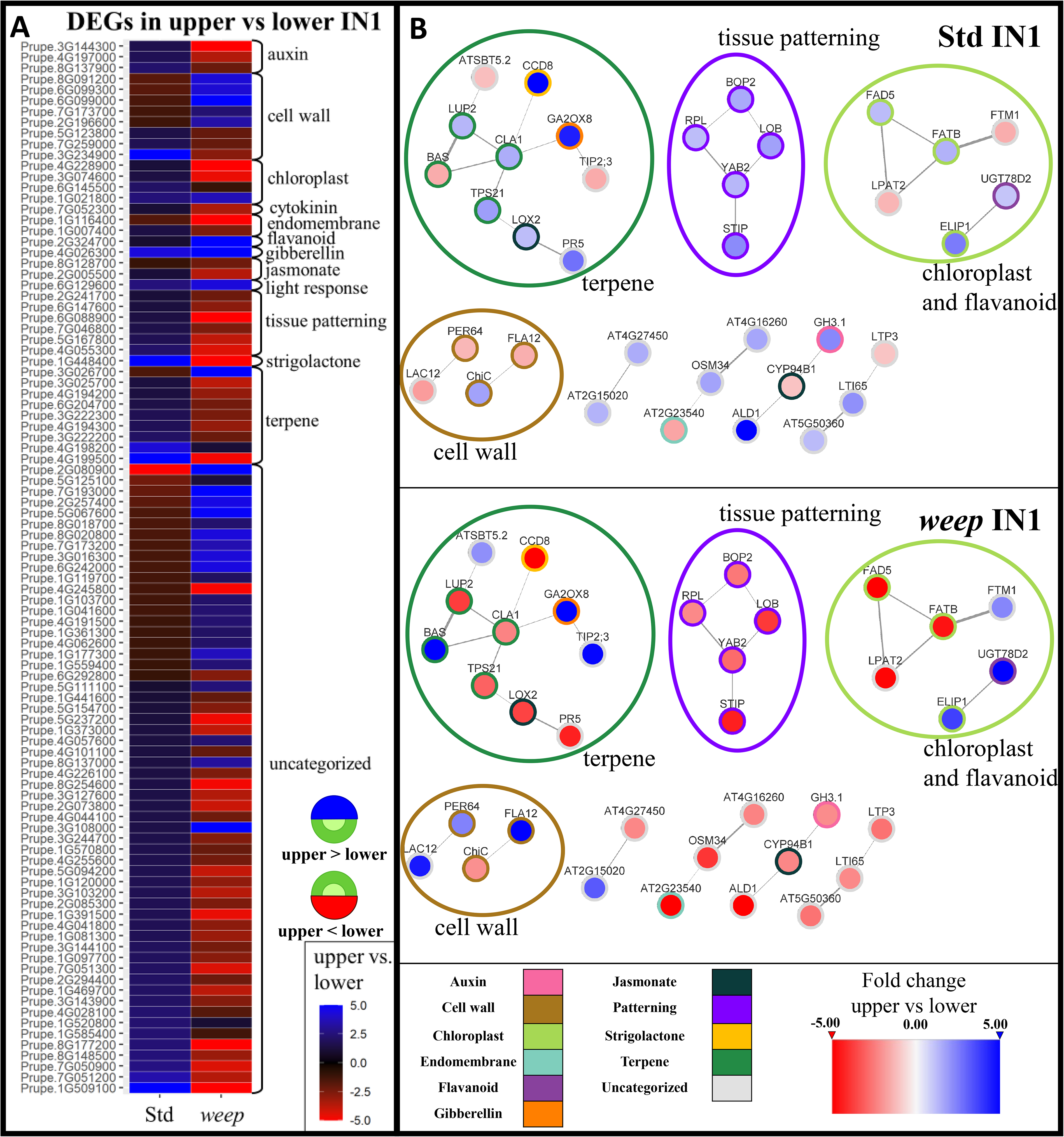
Differentially expressed genes between upper and lower tissues in the first internode (IN1). (A) Heatmap indicates fold changes between the upper and lower sides of shoots from standard and weeping (weep) trees. Red indicates that the expression is higher on the lower side, blue indicates that the expression is higher on the upper side. (B) An interaction network between the genes was created using arabidopsis homologs in STRING. Node color indicates fold change between upper and lower, edge width indicates evidence strength, and node outline indicates functional category assignment.

**Figure 7:**
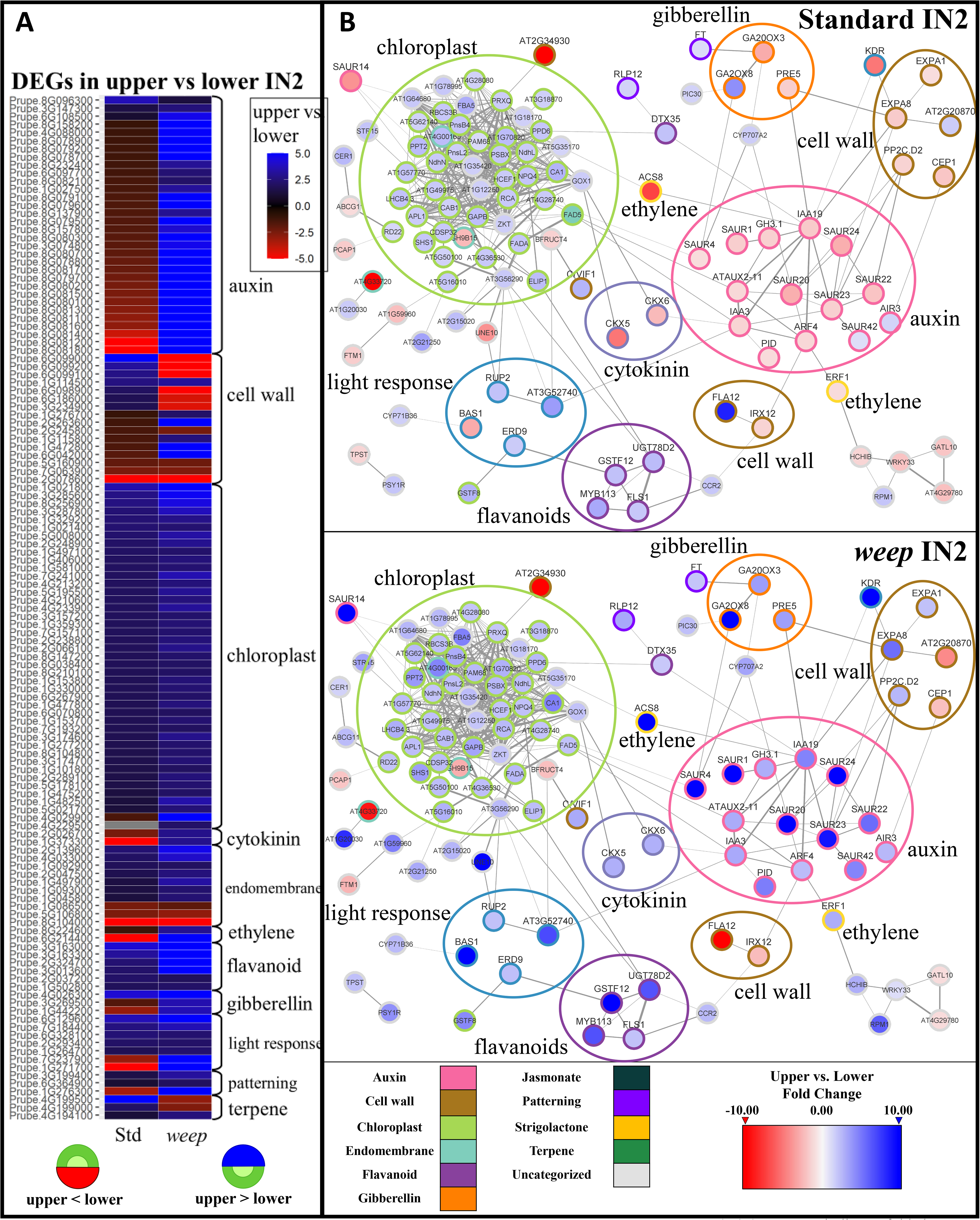
Differentially expressed genes between upper and lower tissues from the second internode (IN2). (A) Heatmap indicates fold changes between the upper and lower. (B) STRING interaction network using arabidopsis homologs. Node color indicates fold change between upper and lower, edge width indicates evidence strength, and node outline indicates functional category assignment.

**Figure 8:**
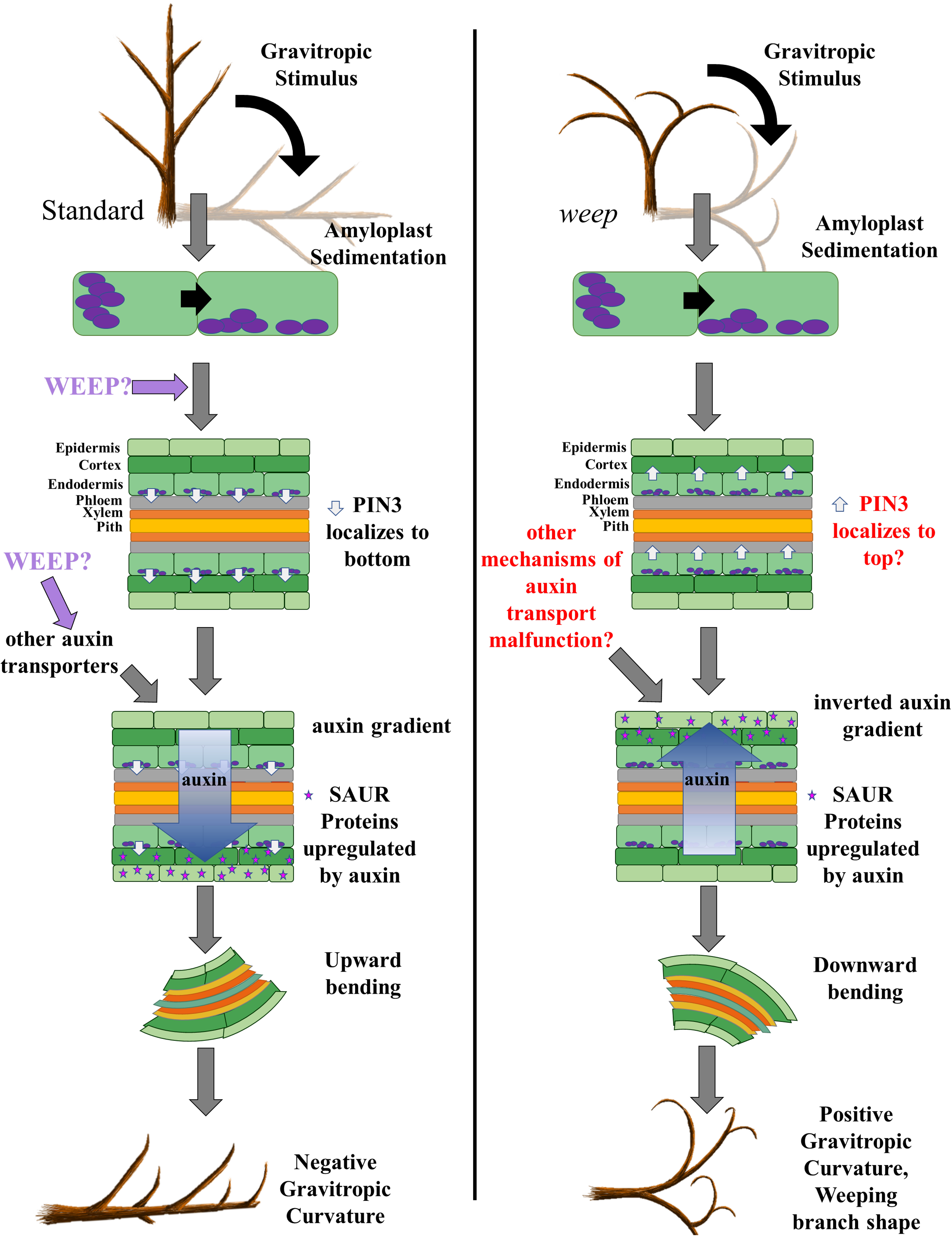
Proposed model of WEEP’s role in shoot tip gravitropism. WEEP acts during gravitropic signal transduction, downstream of amyloplast sedimentation. WEEP is required for formation of the gravitropic auxin gradient, either through localization of PIN proteins or through an alternative auxin transport mechanism. The auxin gradient is required for the shoots’ negative gravitropic bending response. An inverted auxin gradient in the *weep* mutant leads weeping peach shoots to exhibit positive gravitropism.

Using gene annotations and functional descriptions, the genes were also manually categorized into 10 functional groups (i.e., auxin, cell wall, chloroplast, endomembrane, flavonoid, gibberellin, jasmonate, patterning, strigolactone, and terpenes) and one uncategorized gene group (Figures 6 and 7; Supplemental Tables S1 and S2). Interestingly, most of the genes that were differentially expressed between upper and lower tissues in standard peaches were also differentially expressed in weeping peaches, but in an inverted pattern (Figures 6 and 7; Supplemental Figure S6). In other words, genes that were more highly expressed in the upper branch tissues of one genotype were more highly expressed in the lower tissues of the other genotype, and vice versa. To further investigate the GOI, gene correlation networks were created using the arabidopsis homologs of each GOI (Figures 6B and 7B).

Of note for IN1, genes related to tissue patterning, auxin, cell wall development, and terpene biosynthesis and terpene-derived hormones were highly represented and showed inverted expression patterns in the weeping branches (Figure 6B; Supplementary Table S1). The tissue patterning group contains genes related to meristem maintenance, adaxial/abaxial polarity, and phyllotaxy (Prupe.6G088900, homolog of *PRS*; Prupe.4G055300, homolog of *STIMPY (STIP)*; Prupe.2G241700, homolog of *REPLUMLESS (RPL)*; Prupe.6G147600, homolog of *YABBY2* (*YAB2)*; Prupe.5G167800, homolog of *LOB*; and Prupe.7G046800, homolog of *BOP2*). Interestingly, all these genes were expressed only slightly higher in the upper tissues in standard, and much more strongly in the lower tissues in weep, suggesting there might be an inversion of abaxial/adaxial polarity in the *weep* branches. The auxin group included a putative auxin synthesis gene (Prupe.3G144300, homolog of AT4G02610) and two genes for conjugating auxin to amino acids (Prupe.4G197000, homolog of *GH3.6*, and Prupe.8G137900 homolog of *GH3.1*). These genes were also more highly expressed in the upper tissue in standard, and in the lower tissue in *weep*. The cell wall group includes genes involved in lignin metabolism (Prupe.7G173700, *PER64* homolog; Prupe.2G196600, *NST1* homolog), cellulose deposition (Prupe.6G099300 and Prupe.6G099000, *FLA12* homologs), and hemicellulose synthesis (Prupe.5G123800, *CSLG3* homolog). Finally, there was a large group of terpene biosynthesis genes and terpene-related hormones. There are nine differentially expressed terpene biosynthesis genes, including four homologs of the sequiterpene synthase *TPS21* (Prupe.4G194200, Prupe.4G194300, Prupe.4G198200, Prupe.4G199500) and two homologs of the monoterpene synthase *TPS-CIN* (Prupe.3G222200, Prupe.3G222300). Seven of the nine differentially expressed terpene genes are more highly expressed in the upper tissues in standard, and more highly expressed in the lower tissues in *weep*. This pattern is also observed for Prupe.1G448400, a homolog of the strigolactone biosynthesis gene *CCD8* (also known as *MAX4*). In contrast, Prupe.4G026300, which is a homolog of the GA sequestering gene *GA2OX8* is more highly expressed in the upper tissues in both standard and *weep*.

Like IN1, for the IN2 comparison, chloroplast, flavonoid, and many light response genes were more highly expressed on the upper side of the shoot in both standard and *weep* (Figure 7 A, B). This is consistent with expectations, as all of those groups would be expected to be upregulated where there is more light exposure. In contrast, several functional groups were more highly expressed on the lower side of the shoot in standard, but more highly expressed on the upper side of the shoot in *weep*. This pattern is observed in 30 auxin-related genes, including 24 *SAUR* homologs (Figure 7A), a homolog of *PID* (Prupe.4G088000), three homologs of *AUX-IAA* transcriptional regulators (Prupe.1G027500, Prupe.3G074800 and Prupe.8G232400), a homolog of auxin response factor 4 (Prupe.6G097700), and a homolog of GH3.1 (Prupe.8G137900). This pattern is also observed in two ethylene-related genes: a homolog of *ETHYLENE RESPONSE FACTOR 1* (Prupe.8G224600), and a homolog of ACC synthase *ACS8* (Prupe.6G214400). The *ACS8* homolog shows particularly dramatic differential regulation as it is 7-fold more highly expressed in the lower tissue in standard, and 76-fold more highly expressed in the upper in *weep*. Finally, this pattern is observed in two gibberellin-related genes (Prupe.3G269500, Prupe.1G442200), and three terpene synthase 21 homologs (Prupe.4G199500, Prupe.4G199000, Prupe.4G194100).

For the cell wall functional group, there are genes relating to two distinct functions, which show distinct patterns of expression. Four genes are related to cell expansion downstream of auxin, and like the auxin response genes, more highly expressed on the lower side in standard and more highly expressed on the upper side in *weep*. These include three expansins (Prupe.1G276700, Prupe.2G263600, Prupe.6G042000) and a protein phosphatase 2C (Prupe.1G115800). In contrast, four homologs of *FASCICLIN-like arabinogalactan-protein 12* (*FLA12*; Prupe.6G098900, Prupe.6G099000, Prupe.6G099100, Prupe.6G099200) which is a marker of tension wood, are more highly expressed on the upper side in standard and on the lower side in weep.

Because 24 *SAUR* homologs were differentially expressed, and *SAUR* transcript expression is known to be an early auxin response to gravitropism, we further investigated expression patterns throughout the *SAUR* family. Using KEGGORTH terms and *Arabidopsis* homolog descriptions, 76 putative *SAUR* proteins were identified in peach, 43 of which are in a tandem array on chromosome 8. As expected, the majority (54/76) of the putative SAUR proteins in the peach genome are expressed on the lower tissue in standard IN2 (Supplemental Figure S7, Supplemental Table S3). However, the majority (49/76) of the SAUR proteins in *weep* IN2 were expressed in the upper tissues (Supplemental Figure S7), including 40 of the genes upregulated on the lower side in standard. This inverted expression trend also occurred for IN1 tissues, although the difference between expression in upper and lower tissues is more subtle (Supplemental Figure S7). Thus, SAURs overall were more highly expressed on the lower side of the shoot in standard shoots, and on the upper side in *weep* shoots.

### PpeWEEP proteins homo-oligomerize in vitro

Oligomerization is essential for the function of some SAM domain proteins (Denay, 2017). For example, the floral meristem identity protein LFY requires head-to-tail homo-oligomerization to fully access DNA binding regions (Sayou, 2016). To test if the peach WEEP protein (PpeWEEP) homo-oligomerizes, size-exclusion chromatography (SEC) was performed. Heterologously-expressed PpeWEEP fused to 6xHis, maltose binding protein (MBP), and a Strep-Tag, with a predicted weight of 60.3 kDa, was sequentially purified via Ni-NTA and StepTactin columns prior to SEC analysis (Figure 9A). The resulting elution had a prominent peak corresponding to 525 kDa, indicating an average of 8 to 10 monomers per complex, suggesting that most PpeWEEP protein exists as homo-oligomer in solution, similar to LFY (Figure 9B, Supplementary Figure S8) (Sayou et al 2016).

**Figure 9.**
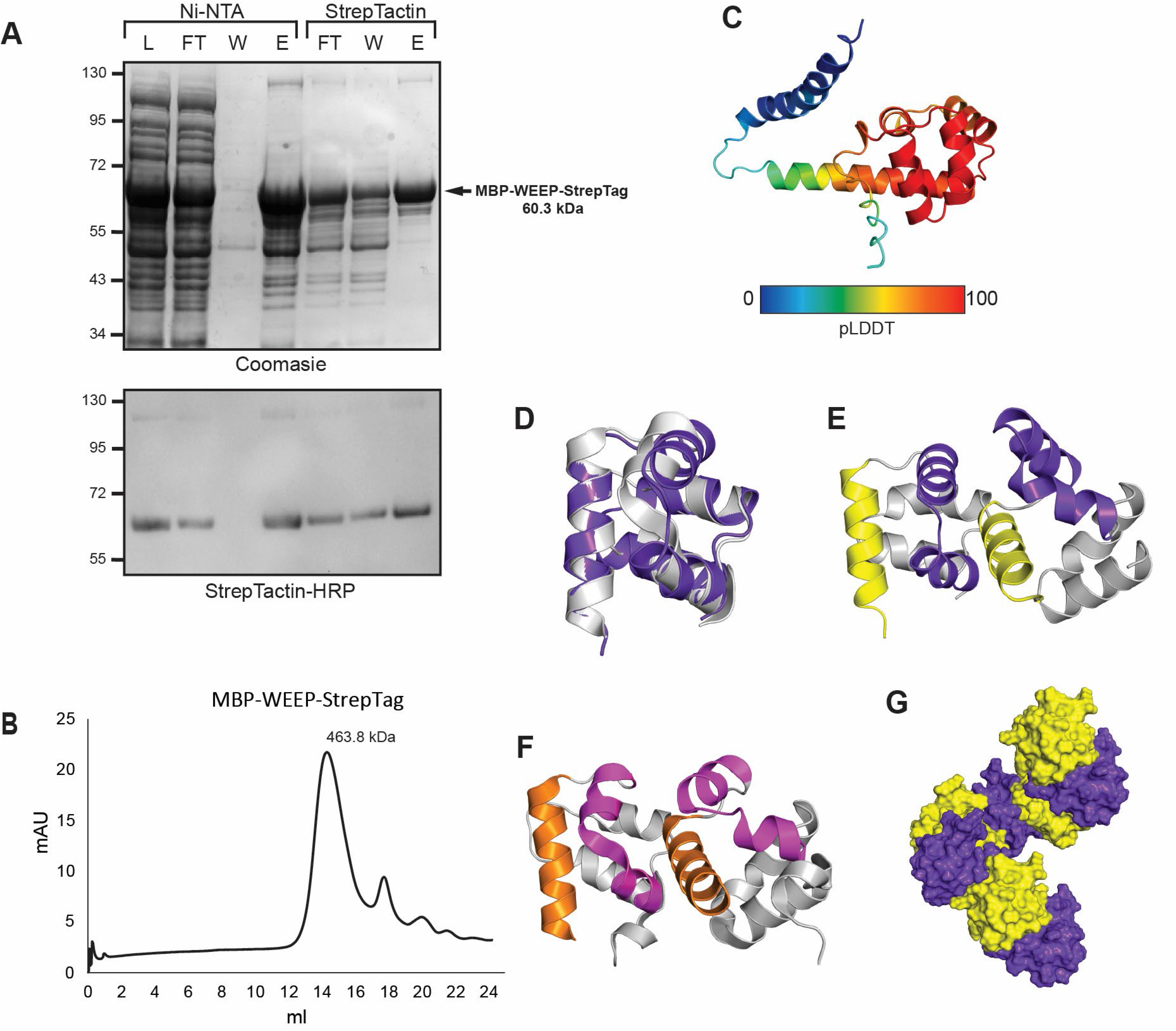
Size exclusion chromatography indicates WEEP proteins homo-oligomerize. (A) SDS-PAGE gels containing flow through (FT), wash (W), and elution (E) samples from sequential Ni-NTA and StrepTactin column purifications of a heterologously expressed 6XHis-MBP-PpeWEEP-StrepTactin fusion protein. Coomassie stained gel (left) indicates total protein. Western blot (right) indicate the presence of the 60.3 kDa fusion protein. (B) SEC chromatograph with a peak at 525 kDa, corresponding to an 8.7mer. (C) AlphaFold2 protein structure predictions for the full length PpeWEEP monomer colored by model confidence (pLDDT) (D) Overlay of WEEP over the top of scm illustrating the similarity between WEEP and scm/Ph. (E-F) AlphaFold2 protein structure prediction for a PpeWEEP SAM domain dimer (E), and the structure of the dimer between the Drosophila SAM domain proteins Polyhomeotic (F) and Sex-comb-on-midleg (Scm) (PDB: 1PK1). (G) Predicted structure of a helical PpeWEEP octomer.

To understand the structure of the PpeWEEP protein, we used AlphaFold2 to predict its structure. AlphaFold2 modeled the structure of the PpeWEEP SAM domain with high confidence, but the C-terminal region with low confidence (Figure 9C; (Bryant et al., 2022). AlphaFold2 multimer modeled the structure of the PpeWEEP SAM domain as a head-to-tail dimer, structurally similar to the Drosophila SAM proteins Polyhomeotic (Ph) and Sex-comb-on-midleg (Scm), despite the low sequence homology between PpeWEEP and Ph/Scm (∼20% identity; Figure 9D-F; Kim et al., 2005). Lastly, PpeWEEP SAM monomers were aligned to the structure of the helical Scm polymer in PyMol to build a model for a PpeWEEP octamer (Figure 9G).

### Weeping peach roots have steeper gravitropic set-point angles and more rapid gravitropic response

In contrast to the absence of a gravitropic response in weeping peach shoots, the roots of *weep* mutants in wheat and barley exhibit both narrower gravitropic set-point angles and a more rapid root positive gravitropic response (Kirschner et al., 2021). Therefore, we assessed weeping peach root set-point angle under normal growth conditions and performed a time-course following 90-degree reorientations to measure the timing and extent of root gravitropic response.

Freshly germinated standard and weeping peach seeds from the same population were planted in rhizotrons, and the resulting seedlings were grown for ten weeks to observe their natural root architecture. The peach *weep* mutant exhibited a dramatically different root architecture than the standard peaches (Figure 10A-C). Driving this change, weeping peach roots have significantly steeper lateral root emergence and tip angles (Figure 10D and E), which leads to a smaller convex hull (area of root exploration; Figure 10F). In addition, the number of secondary and tertiary lateral roots was moderately increased in *weep* (Figure 10G). Lastly, multiple weeping tree secondary roots grew straight downwards and were often co-dominant with the primary root (Fig 10A and B). Examination of the root system at the end of the experiment after soil removal confirmed that these were true secondary roots, rather than seminal roots emerging from above the radical (Figure 10C).

**Figure 10.**
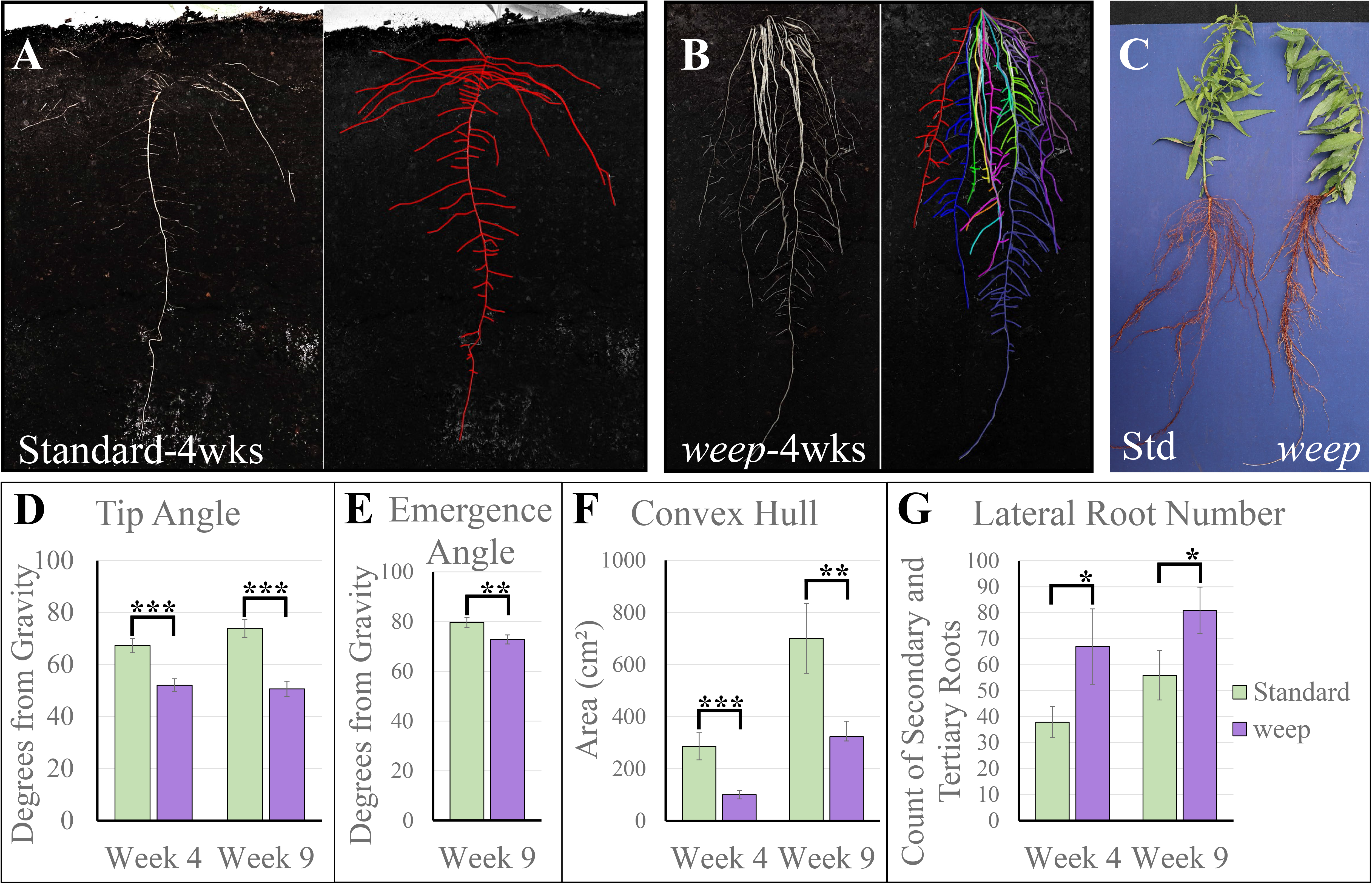
Root architecture in standard and weeping peaches. Peach seedlings were grown in rhizoboxes, imaged, and the root system was traced in RootNav (A, B). Different colors indicate separate primary roots, or secondary roots with lateral tertiary roots. RootNav was used to measure tip angle (D), emergence angle (E), convex hull (F), and lateral root number (G). At the end of the experiment, dirt was removed, and the entire root system was photographed (C). Pairwise comparisons done with t-tests. Error bars show standard error. * indicates significantly different at α=0.10; ** indicates significantly different at α=0.05; *** indicates significantly different at α=0.01.

Next, we assessed gravitropic response of actively-growing root-pruned seedlings following a 90-degree reorientation (Figure 11A). The seedlings had undergone root pruning prior to planting to increase the root numbers, as pruning promotes vigorous lateral root growth. Before reorientation, vertically oriented roots in each rhizotron were selected, and root tip angle was measured from the initial trajectory of these roots. In response to 90-degree reorientations, weeping tree roots exhibited a faster gravitropic response than standard, with a significantly greater root tip angle at two days after reorientation (Figure 11B). The two genotypes had the most similar response at four days after rotation (Figure 11B). Between four and six days, the standard root tip returned to a more horizontal trajectory, leading *weep* to once again have a significantly wider angle at six days after reorientation (Figure 11B). This suggests that the set-point angle of weep roots is decreased.

**Figure 11.**
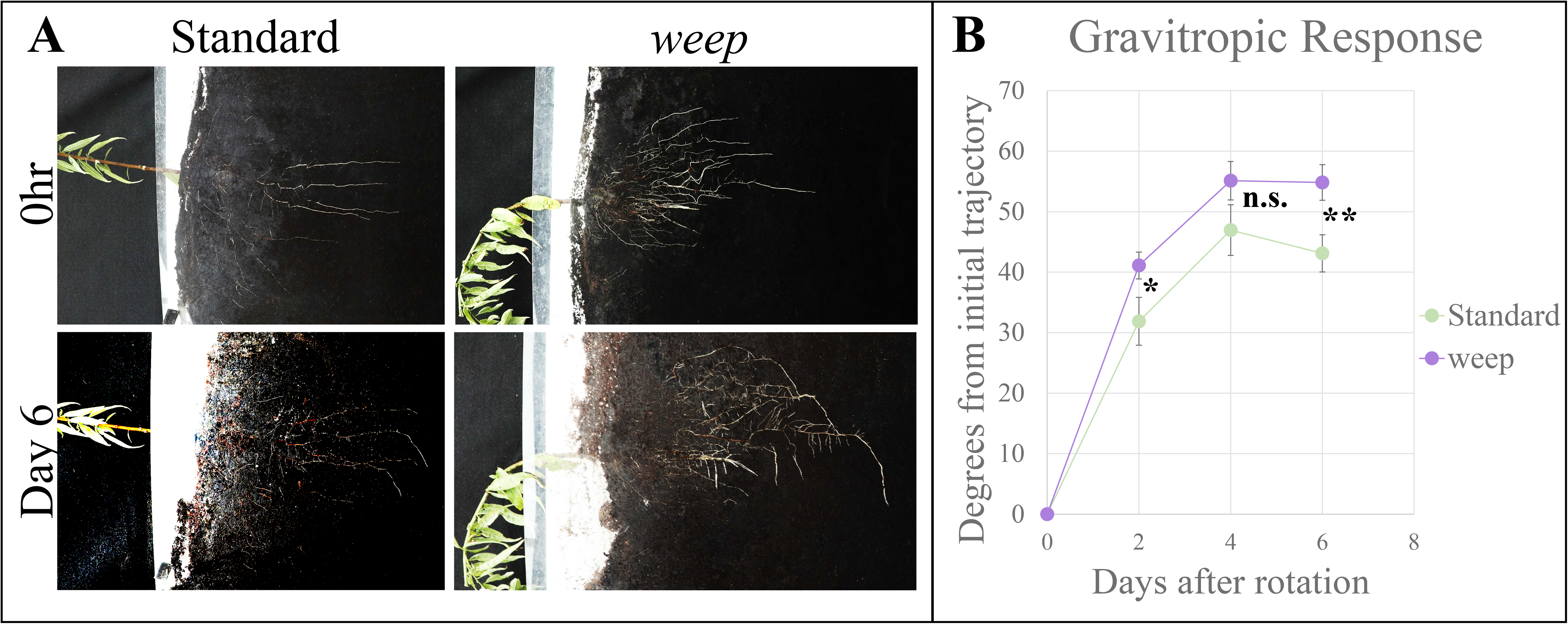
Root gravitropic response. Standard and weep seedlings were rotated 90° and photographed over the course of 6 days to assess root gravitropic response (A). The root tip angle from initial trajectory was measured at 2,4, and 6 days (B). Pairwise comparisons done with t-tests. Error bars show standard error. * indicates significantly different at α=0.10; ** indicates significantly different at α=0.05; *** indicates significantly different at α=0.01.

## Discussion

### WEEP directs the formation of the asymmetric auxin gradient needed for shoot gravitropism

Unlike other weeping *Prunus* trees, weeping peaches do not have altered branch anatomy or decreases in branch stiffness. Changes in the proportion of xylem in the shoot tip are transient, disappearing by the end of the growth season. No significant differences in cellulose and lignin content or localization were detected between weeping and standard peaches. Furthermore, the flexural stiffness and other biomechanical properties of young shoots are not different between standard and *weep.* Collectively, these data suggest that weeping peach branches grow downwards and fail to reorient upwards not due to failure under self-loading, but because of an alteration in gravitropic response.

Several lines of evidence indicate that this altered shoot gravitropic response is due to an inversion of the auxin gradient in weeping peach trees, which leads to positive (rootward) gravitropic bending. To pinpoint the role of *WEEP* in the gravitropic pathway, we assessed gravitropic perception, gravitropic signal transduction, and gravitropic response in the *weep* peach mutant. The normal endodermis development and amyloplast sedimentation in peach *weep* shoot tips suggest that *weep* shoots have normal gravitropic perception (Figure 4A and B). This agrees with the previous finding that amyloplast sedimentation is normal in barley *egt2* mutant root columella cells (Kirschner et al., 2021). Next, we demonstrated *weep* shoots showed a normal bending response to unilateral auxin application. Notably, auxin application to the bottom of *weep* shoot tips could partially reverse the *weep* phenotype, causing them to grow upwards (Figure 5A). Conversely, application of auxin to the top of standard shoots phenocopied *weep*’s downward bending of the shoot tip (Figure 5B). Normal auxin response has also been observed in barley *egt2* roots, where auxin inhibits elongation and gravitropic bending in both wild type and *egt2* (Kirschner et al., 2021). As weeping peach trees are not impaired in either gravitropic perception or response, our results suggest the WEEP protein functions in gravitropic signal transduction upstream of the auxin gradient formation.

RNA sequencing from upper and lower shoot tip tissues from standard and weeping peach trees revealed inverted localization of early-auxin response gene expression in the *weep* mutant. Early auxin response genes provide a good marker for auxin localization, as they are rapidly upregulated by auxin (often within a few minutes). They are generally categorized in three classes: GH3s, AUX/IAAs, and SAURs, with1-amino-cyclopropane-1-carboxylate synthases (ACSs) sometimes included as a fourth class (Grossmann, 2010; Ren and Gray, 2015; Pei et al., 2019). The expression of these genes results in a highly interconnected network which immediately acts downstream of auxin to stimulate shoot elongation (Figure 7B). Genes in all four classes were differentially regulated between upper and lower tissues in IN2 in both *weep* and standard (Figure 7A and B, Suppl. Figure S2). Specifically, they all exhibited higher expression on the lower side of standard peach shoots, but higher expression on the upper side of the weeping shoots (Figure 8). The differential expression pattern in standard shoots suggests the mechanism behind the regulation of upward branch orientations aligns with the Cholodny-Went gravitropic response pathway: higher auxin concentrations on lower (abaxial) branch tip tissues likely promote upward growth trajectories by promoting cell elongation. Accordingly, the flipped expression pattern of the early auxin response genes in the weeping shoot tips suggests that the downward growth trajectory is due to a higher auxin concentration in the upper tissues.

Of particular note is the large number of *SAURs* upregulated on the lower side of IN2 in standard and the upper side in *weep* (40 out of the 76 putative SAUR genes we identified in peach, Supplemental Figure S7, Supplemental Table S3). *SAUR* genes have previously been identified as upregulated on the lower side of shoots during gravitropism in soybean (*Glycine max*) and *Arabidopsis* (Ren and Gray, 2015; Wang et al., 2020). Most *SAURs* are very rapidly induced by auxin (often within 5 minutes) through the canonical SCF^TIR1/AFB^ signaling pathway (Du et al., 2020). Auxin transport is necessary for asymmetric expression of *SAURs* during gravitropism, suggesting that this asymmetric expression is entirely generated by the auxin gradient, and there is no known alternative pathway for localization of *SAUR* expression (Du et al., 2020; Wang et al., 2020). Thus, *SAUR* expression is a reliable indicator of auxin localization. The upregulation of SAURS and representatives of all the other classes of early auxin response genes on the upper side of weep shoots leaves little doubt that the auxin gradient is inverted in *weep,* with higher levels of auxin on the top of gravitropically stimulated *weep* stems.

Furthermore, *SAUR* expression is both necessary and sufficient for cell elongation. During shoot gravitropism responses, SAUR proteins on the lower side of shoots inhibit PP2C-D phosphatases from dephosphorylating H^+^-ATPases (Spartz et al., 2014; Du et al., 2020). The phosphorylated H^+^-ATPases are activated and pump protons out of the cell, hyperpolarizing the cell membrane. That hyperpolarization leads to water uptake by the cell and a resultant increase in turgor pressure, as well as apoplast acidification, which activates expansins and other cell wall remodeling enzymes (Spartz et al., 2014; Du et al., 2020).Collectively, these events lead to cell elongation on the bottom side of gravitropically stimulated shoots, reorienting the shoots upward. *SAUR* expression is necessary for auxin-mediated cell elongation as *saur* knockouts show dramatically reduced gravitropic responses and *pp2c.d2* mutants that are insensitive to *SAUR* regulation are also insensitive to auxin. *SAUR* expression is also sufficient to independently stimulate cell expansion. Plants with constitutive overexpression of *SAURs* exhibit enhanced cell expansion (Du et al., 2020; Wang et al., 2020).

In accord with the expected effects of *SAUR* expression on cell elongation, in standard peach shoot tips homologs of *Arabidopsis* cell expansion promoting genes *PACLOBUTRAZOL-RESISTANCE5* (*PRE5*), *EXPANSIN A1* (*EXPA1*), and *EXPA8* (Prupe.3G269500, Prupe.1G276700, and Prupe.2G263600) are more highly expressed on the lower side of the shoot (Figure 7A; Cosgrove, 2015; Shin et al., 2019). In contrast, in *weep* shoots, these genes’ expression is inverted, being more highly expressed on the upper side of the shoot. Although we were unable to quantify differences in cell length from shoot tip sections, our expression data strongly suggest that increased auxin levels in the upper side of shoot tips lead to increased cell elongation, which would result in the downward growth phenotype in weeping peach. The decrease in fiber cell size in weeping peach branches also suggests WEEP plays a role in cell elongation, however slight, although significant upper vs. lower tissue differences were not detectible from our hand dissections. Reduced expression of genes related to cell elongation was also observed in the root elongation zone of *egt2* mutants (Kirschner et al., 2021). Because *egt2* roots showed normal inhibition of growth when auxin was exogenously applied, Kirschner et al. concluded that *egt2* likely acted independently of auxin in control of cell elongation. Our auxin application data also show a normal auxin response in *weep* peach. However, the role of WEEP in controlling endogenous auxin localization within gravitropically stimulated tissues provides a parsimonious explanation for the alterations in cell elongation. Indeed, this hypothesis is more in accord with Kirschner’s results than a direct role for *WEEP* in cell elongation, as *egt2* mutants in barley did not show altered root length (Kirschner et al., 2021).

In addition to the canonical effects of auxin localization on cell elongation, auxin localization is also essential for lateral organ boundaries and adaxial/abaxial patterning. Local auxin maxima are required for lateral organ initiation (Heisler and Byrne, 2020). While multiple theories have been proposed for determination of adaxial/abaxial polarity, several lines of evidence (including auxin applications, microdissections, application of polar auxin transport inhibitors, and confocal microscopy of PIN1 and auxin reporters) suggest that polar auxin transport and auxin localization is crucial to maintaining polarity (Shi et al., 2017; Heisler and Byrne, 2020; Burian et al., 2022; Wang et al., 2022). In accord with this data and the hypothesized role for WEEP in auxin localization, many tissue patterning genes showed inverted expression in *weep* IN1. Of these, five were homologs of Arabidopsis genes which formed a small gene network (Figure 6B). Among these is YAB2, which is expressed on the abaxial side of lateral organs and is essential for organization of the shoot apical meristem and adaxial/abaxial differentiation (Stahle et al., 2009). Also in the group is BOP2, which is essential for proximal/distal patterning in leaves, radial patterning of flowers, promotes adaxial development, and upregulates LOB, which is essential for establishing the boundary between the meristem and lateral organs (Hepworth et al., 2005; Žádníková and Simon, 2014). Another member is *RPL,* which represses *AGAMOUS* and is involved in floral whorl differentiation and specification of phyllotaxy (Bao et al., 2004; Gish, 2013). Finally, *STIP* controls cell fate in the meristem, repressing *WUSCHEL,* promoting cell division, and preventing differentiation (Wu et al., 2005). Together, the altered localization of these genes in *weep* suggests that the mislocalization of auxin may be affecting meristem patterning.

### The role of WEEP in auxin gradient formation may be essential to localizing tension wood formation

The inversion in auxin gradient and tissue polarity may also result in a mislocalization of tension wood in weeping peaches. Although cellulose and lignin staining and extractions did not indicate the presence of tension wood in either upper or lower regions of standard or *weep* branches (Figure 2C, D), our RNAseq data did show differential expression of genes associated with tension wood.

High *FLA* expression is associated with tension wood formation in *Eucalyptus,* willow (*Salix*), and poplar (*Populus*; Qiu et al., 2008; Gritsch et al., 2015; Wang et al., 2017). The *Arabidopsis* homolog *AtFLA12* is a crucial trigger of cellulose microfibril deposition in stem vascular tissues (MacMillan et al., 2010). Double mutants of *Atfla12* and related gene *Atfla11* have even further decreases in cellulose content and higher microfibril angles (MacMillan et al., 2010). Thus, FLAs appear to induce both the higher cellulose content and lower microfibril angles, which are characteristic of tension wood. Four homologs of *AtFLA12* were upregulated on the upper side of standard peach shoots. This was consistent with the expression pattern of the eucalyptus homologs of *AtFLA12, EgrFLA1* and *EgrFLA2,* and indicates tension wood formation in the upper portion standard peach branches (Qiu et al., 2008). In contrast, all four peach homologs were upregulated on the lower side of weeping shoots. Accordingly, the downward curvature of weeping shoots may be associated with the formation of tension wood on the lower side of *weep* branches.

Further research is needed to understand how the formation of tension wood is connected to the gravitropic auxin gradient in young shoots which are undergoing both elongation and radial growth through xylem formation. Tension wood formation is associated with upregulation of auxin, ethylene, and gibberellic acid (GA) signaling, and FLAs are upregulated by GA in poplar (Gerttula et al., 2015; Wang et al., 2017). Yet, in young shoots, such as we assessed here, auxin, ethylene, and GA are all coordinately upregulated on the lower side of the shoot, opposite the location of tension wood. Indeed, a homolog of the rate-limiting enzyme for ethylene biosynthesis, 1-amino-cyclopropane-1-carboxylate synthase 8 (ACS8) was upregulated 7-fold on the lower side in standard, but 76-fold on the upper side in weep. Similarly, the homolog of GA20OX3, a GA synthesis enzyme, is 3-fold upregulated on the lower side in standard and 4-fold upregulated on the upper side in *weep.* One possible solution is to look at auxin flux not at the epidermal cells (which are involved during primary growth in shoot elongation) but at the vascular cambium, where xylem formation is taking place (Gerttula et al., 2015). Under this paradigm, during gravitropism auxin is moving away from the epidermis in the upper part of the branch, but toward the vascular cambium, stimulating xylem development and tension wood formation (Gerttula et al., 2015). On the lower part of the branch, auxin is moving away from the vascular cambium, and toward the epidermis, promoting cell elongation. In *weep* the auxin flux would be toward the vascular cambium on the lower side, and toward the epidermis on the upper side, which would then be consistent with the formation of tension wood on the lower side of the branch and increased cell elongation on the upper side.

### WEEP contributes to the regulation of antigravitropic offsets by promoting polar auxin transport

In agreement with studies in other species (Kirschner et al., 2021; Johnson et al., 2022), we found the weeping peach root system displays decreased lateral root angles, a narrower convex hull, a faster positive gravitropic response, and a more vertical setpoint angle. This suggests that *WEEP* normally promotes negative gravitropism and subsequent maintenance of upward growth in peach roots, as well as in shoots.

As in shoots, auxin gradients are required for gravitropic responses in roots and auxin concentrations are higher in the lower tissues. But, in contrast to shoots, high auxin concentrations inhibit cell elongation in roots. This response is very dosage dependent, as low levels of auxin in the upper tissues of gravistimulated roots stimulate the elongation that promotes downward growth (Du et al., 2020). Both shoot and root angles are narrower (more vertical) in mutants with higher auxin or auxin response, while mutants with lower auxin or auxin response show wider lateral angles (Roychoudhry et al., 2013).

However, lateral shoots do not grow vertically up, nor do lateral roots grow vertically down. Rather, lateral branches and roots grow at and maintain a genetically-encoded angle known as the gravitropic set-point angle, which must be maintained by an anti-gravitropic offset (AGO) (Roychoudhry et al., 2013). The presence of this anti-gravitropic offset has been confirmed by clinorotation and microgravity experiments where lateral organs show marked outward growth in the absence of gravitropism, but the mechanism is unknown (Roychoudhry et al., 2013). Crucially, auxin is required for this response, as the outward growth is inhibited in the absence of auxin transport (Roychoudhry et al., 2013). Thus, auxin localization appears to control both the gravitropic response and the anti-gravitropic offset, and auxin homeostasis is likely crucial to setting the balance between the two.

We hypothesize that the WEEP protein functions in both shoot and root gravitropism and AGO pathways by modifying polar auxin transport to create auxin gradients. More specifically, we suggest that the WEEP protein promotes the upward shoot growth by promoting the transport of auxin to lower tissues of gravistimulated vertical shoots and upward orientated lateral branches. And in roots, we suggest that WEEP decreases gravitropic responses and promotes the maintenance of non-vertical downward lateral root growth through a homeostasis mechanism, which decreases the auxin gradient between upper and lower tissues by promoting the movement of auxin to upper side of lateral roots. Essentially, the WEEP protein may be a key player in the maintenance of the anti-gravitropic offset (AGO) associated with non-vertical growth.

Further experimentation is needed to investigate the mechanism by which *WEEP* controls auxin homeostasis and transport during gravity perception. However, it is likely that WEEP directly or indirectly modifies the localization of the PIN3 auxin efflux proteins in the plasma membrane of statocytes. PIN protein localization is controlled by endosomal trafficking (Rakusová et al., 2015; Zhang et al., 2020). The endosomal trafficking is regulated by the PINOID kinase, which phosphorylates PIN proteins (Kleine-Vehn et al., 2009). SAM domain proteins are known to be involved in vesicle trafficking. Arabidopsis SAM1 interacts with four vesicle trafficking proteins (Wang et al., 2011) and STIM1, a human SAM protein, interacts with microtubules (Grigoriev et al., 2008). *WEEP* homologs have been localized to the plasma membrane, cytoplasm, and nucleus (Kee et al., 2009; Guo et al., 2023) Arabidopsis WEEP protein (AtSAM5) interacts with the nuclear- and plasma-membrane-localized kinase CPK13, suggesting a potential role in regulating PIN phosphorylation at the plasma membrane (Denay et al., 2017). Alternatively, WEEP may act in transcriptional regulation of vesicle trafficking. The mutant of the barley *WEEP* homolog, *egt2,* showed upregulation of exocyst complex component 7 (EXOCYST70A3), which is known to be involved the distribution of PIN4 and root gravitropic responses (Ogura et al., 2019; Kirschner et al., 2021).

Self-oligomerization of WEEP proteins through their SAM domain may also be key to its function. Oligomerization through these domains is essential for the subcellular localization and activity of some SAM proteins (Denay et al., 2017). Oligomerization through the SAM domain is essential to the action of LFY as a transcriptional regulator (Sayou et al., 2016). Regardless of whether WEEP acts at the plasma membrane, the nucleus, or both, the absence of any known protein motifs besides the SAM domain suggests that WEEP may serve as a scaffold or structural protein regulating the formation of protein complexes.

## Conclusion

Collectively, our peach shoot and root data provide exciting additions to our understanding of how plants respond to gravity and maintain non-vertical growth orientations. *WEEP* promotes negative gravitropism in both shoots and roots. In peach shoots, which are normally negatively gravitropic, mutations in *WEEP* lead to a positively gravitropic phenotype, where the shoots grow towards the ground. The transcriptomic evidence we present here suggests that *WEEP* is necessary for normal establishment of a gravitropic auxin gradient in peach shoots, and that gradient is inverted in *weep* peach mutant shoot tips. Further research is needed to elucidate whether the WEEP protein plays a role in auxin localization in shoots by promoting PIN3 protein localization or whether it acts through an orthogonal mechanism. The absence of a clear shoot phenotype in barley, wheat, and arabidopsis *weep* mutants suggests the processes of lignification and/or altered tension wood localization are necessary to “freeze” the curvature in time and produce a visible phenotype.

## Materials and Methods

### Plant Material

Experiments used 1- to 4-year-old standard and weeping peach trees grafted on Halford rootstock, unless otherwise noted. The peach scions originated from the segregating F2 population used to map the *weep* gene (Hollender et al., 2018). Trees were grown in pots ranging in size from 1- to 15-gallons in standard greenhouse conditions, with supplemental lighting to maintain an approximately 16-hour photoperiod. Dormancy requirements were met by placing the trees once or twice a year into a 4°C dark cold room for at least six weeks at a time.

### Sectioning for stem anatomy

Actively growing shoot tips and dormant first-year branches with outward growth orientations were hand-sectioned with a double-edged razor blade. Actively growing shoot tips were sectioned at approximately 6cm below the shoot apical meristem from three shoot-tips per tree with four trees for each genotype. Dormant branches were sectioned at about 6 cm below the tip from three shoot-tips per tree, with four trees for *weep,* and three trees for standard. Three clean sections were taken from each shoot tip. The sections were stained with 1 mg/ml Toluidine blue freshly diluted to 50% in water and photographed using a Nikon SMZ800N dissecting microscope with a Nikon DS-Fi3 camera.

Each of the three sections was measured in ImageJ using the polygon measuring tool. The pith was measured by tracing around the parenchyma cells. Xylem was measured around the outside of the rays where toluidine staining was clearly visible. The total area of the section was measured tracing around the outermost cells of the epidermis, which were clearly stained.

Data were modeled using the lme4 library in R (v. 4.2.1) with blocking by tree and by shoot tip (response= mean + Genotype + Tree:Genotype + Shoot_Tip:Tree:Genotype + error, where response is the pith area or xylem area, Genotype is the effect of genotype, Tree is the effect of the individual tree, nested within genotype, and Shoot_Tip is the effect of the shoot tip, nested within tree and genotype. The effects of tree and shoot tip were treated as random variables. Data were assessed for normality and equal variance, then pairwise comparisons between genotypes were performed using emmeans to apply two-tailed t-tests assuming equal variances.

### Wood composition

Segments, approximately 1.5 cm in length, were taken from two-year-old wood from weeping and standard peach branches from the original mapping population at the USDA Appalachian Fruit Research Station (Kearnysville, WV). At collection time the ‘upper’ and ‘lower’ (adaxial and abaxial) sides of the branches with respect to gravity were marked. These segments were then bisected longitudinally, the bark and pith were removed, as well as some of the edge regions orthogonal to the gravity vector. This produced roughly trapezoidal shaped fragments of wood representing the woody growth of the top or bottom of a branch. These were cut into roughly 0.5 x 0.5 cm pieces with a sharp razor, extracted in series with 70% EtOH and 100% acetone, each overnight. Then, samples were dried and ball milled into a fine powder. The crystalline cellulose content, non-cellulosic polysaccharide content, acetyl-bromide soluble lignin content, and lignin monomer content were then assessed as described previously (Foster et al., 2010a; Foster et al., 2010b). For each measurement, three technical replicates were performed for four biological replicates per sample type.

Data were modeled using the lme4 library in R (v. 4.2.1) with blocking by sample (response= mean + Genotype + Side + Genotype*Side + sample:Genotype + error where response is the cell wall polymer, Genotype is the effect of genotype, Side is the effect of top or bottom, Genotype*Side is their interaction, and sample is a random variable nested within genotype. Data were assessed for normality and equal variance, then all pairwise comparisons were performed using emmeans to apply Tukey tests assuming equal variances. Paired t-tests between top and bottom within a genotype were performed in Excel.

### Fiber cell measurements

Wood samples from the top and bottom of dormant two-year old branches were dissected as described for wood composition and then cut into approximately 1 x 0.2 cm ‘match sticks.’ Samples were then macerated by the addition of 10 mL of Franklin’s solution (50% acetic acid, 4% hydrogen peroxide) and incubation at 60°C for 2-3 days, followed by heating in a boiling water bath for 10 minutes. Franklin’s solution was then removed, and the samples were then washed at least three times with 10 mL H_2_O. Vigorous mixing by vortex was then able to fully break apart the fiber cells (Chaffey, 2002; Franklin, 1945). Fiber cells were stained with 0.01% Safranin O in H_2_O, mounted on microscope slides, and 12 pictures per sample were taken. The length of all full-length fiber cells visible in each picture were measured with NIS Elements (Nikon) and the data were exported to excel. Statistical analysis was performed as described for wood composition, except ANOVA was significant, so t-tests were used.

### Wood material properties

Three-point bending tests were conducted using a universal testing machine (Instron^©^ Model 4202, Instron Corporation; Kern et al., 2005). Tests were performed on four shoot tips per tree, for three standard and four *weep* trees). Shoot tips were selected to have 2-3 mm diameter. Approximately 10 cm segments were used, starting 3 cm beneath the shoot apex, as the shoot’s stiffness varied widely in the first 3 cm, but were reasonably consistent from 3 cm on down. The top of the shoot was placed downward so that the top of the shoot was under tension and bottom under compression, as it would be under self-loading. The total span length was 8cm, with the cross-head centered. The load was applied until failure using a 50 N load cell and a cross-head constant speed of 20.00 mm/min.

Modulus of rupture (MOR) was calculated as MOR= (½*F*max)(*a*)(R_2_)/*I*, where “*F*max” is the maximum force recorded by the Instron, “*a*” is the distance between the post and load, “R_2_” is the radius of the sample perpendicular to the direction of the load, and *I* is the second moment of inertia. *I* was calculated for an elliptical cross-section as *I*=(π)(R_1_) ^3^ (R_2_) /4, where “R_1_” is the radius in the direction of the load. Flexural stiffness (*EI*) was calculated using the linear (elastic) portion of the stress/strain curve. EI=(F/V)(a^2^/12)(3L-4a), where F/V is the slope of the stress/strain curve, and *L* is the total span length. Modulus of elasticity (MOE) was calculated as MOE=*EI*/*I*.

### Amyloplast staining of vibratome sectioned tissues

An area about 1 cm below the apex was cut into approximately 50 μm radial sections, using a Vibratome Series 1000 sectioning system. These were stained with 100% Lugol’s iodine solution for several seconds, de-stained in water, and immediately observed under a Nikon ECLIPSE Ni upright microscope with a Nikon DS-Fi3 camera.

### Tissue Collection, fixation, and embedding for sectioning for amyloplast sedimentation analysis

Shoot tips were harvested and cut to separate apices from the 0.5 cm long stem section below them. Cuts were done so that the orientation of the 0.5 cm stem samples relative to the shoot tip could be determined, and tissues were marked with a permanent marker to denote the direction of the gravity vector. Next, they were immediately submerged in ice-cold fixative (2.5% glutaraldehyde / 2.5% formaldehyde / 0.1 M sodium cacodylate buffer, pH 7.2). Samples were vacuum infiltrated for 30 min at room temperature and then the fixative was replaced with fresh solution. This process was repeated once before storing the samples in fixative at 4°C until embedding. Prior to dehydration and embedding, drawings of each sample were done to preserve the gravity vector information (this was possible because the samples were sufficiently asymmetrical). At the start of the embedding process, fixative was removed, and samples were washed three times with 0.1 M sodium cacodylate buffer (pH 7.2) before dehydrating them with increasing concentrations of acetone (30%, 50%, 70%, 80%, 90%, 100%, 100%, 100%). Samples were vacuum infiltrated for 30 minutes at each concentration then gently shaken for 30 minutes in fresh solution, and then shaken overnight in 100% acetone. Next, tissues were gradually infiltrated with increasing concentrations of a modified low-viscosity epoxy resin (10 g ERL-4221 (vinyl cyclohexene dioxide), 6 g D.E.R. 736 (diglycidyl ether of polypropylene glycol), 26 g NSA (nonenyl succinic anhydride), and 0.2 g DMAE (dimethylaminoethanol); Spurr, 1969). Spurr:acetone solution ratios in order were 1:3, 1:3, 1:2, 1:1, 2:1, 3:1, with four 100% Spurr incubations. Each concentration was carried out for one full day. In the morning, samples were vacuum infiltrated for 1 hour in the solution for that day and then shaken in fresh solution for at least 5-12 hours. This process was repeated in the evening. At the end of the infiltration sequence, tissues were positioned in block molds with respect to the gravity vector and polymerized in Spurr resin at 60 °C for 48-72 hours.

### Sectioning, staining, and imaging

Tissues were sectioned with a diamond knife and an RMC PTXL ultramicrotome in the Center for Advanced Microscopy (Michigan State University, East Lansing, MI). Three samples were sectioned per genotype and tissue type (12 samples total) at 1 μm thickness. To control for location in the tissue, each sample was sectioned to the approximate center, where the pith was the widest. After adhering the plastic sections to glass slides using a hot plate, they were stained with a 25% toluidine blue and basic fuchsin solution (Electron Microscopy Sciences Cat #14950). Sections were imaged with a Nikon Eclipse Ni compound microscope with a Nikon DS-Fi3 color camera and NIS-Elements BR 4.60.00 software (Nikon).

### Auxin applications

Auxin was applied to young shoots (1.5-5cm long) on greenhouse-grown trees. 1% w/w IAA in lanolin (100 µl of 100 mg/ml IAA in 1 M KOH per 1 g of lanolin) or lanolin control (100 µl 1 M KOH per 1 g of lanolin) was applied unilaterally to either the top or the bottom of the shoot. Three replicates of each treatment (IAA top, control top, high IAA bottom, and control bottom) were performed per tree, for two trees per genotype. Shoots were photographed at the beginning of the experiment, and at 1, 2, 3, and 7 days after application.

For detached shoots, growing shoot tips were collected in the summer of 2020 from a field planting in Clarksville, MI from three-year-old trees of *weep* and the standard cultivar ‘Bounty’. Three replicates were performed per treatment, and the experiment was repeated twice. After removing the leaves, twigs were placed upright in distilled water under 16-hour (first replicate) or 24-hour (second replicate) light at room temperature. The twigs were first allowed to acclimate for 12+ hours to avoid gravitropic curvature responses, then 1% auxin/lanolin paste (100 µl of 100 mg auxin/ml 1 M KOH stock solution /g lanolin) or control paste (100 µl 1 M KOH stock/g lanolin) was applied to one side of each twig. Photographs of the twigs were taken every 5 minutes for 5 days using a Canon EOS M5 camera.

### Hormone analysis

We selected actively growing branches with an outward orientation (roughly perpendicular to the gravity vector). Two sections were collected: internode 1 (IN1; from the shoot apex to the base of the first elongated internode) and internode 2 (IN2; the next elongated internode). Each section was bisected into top and bottom with respect to the gravity vector, and the samples were immediately flash-frozen in liquid nitrogen. For hormone analysis, each sample type was pooled from 6-8 shoot tips from the same tree. Three standard trees and four *weep* trees grown under the same greenhouse conditions were used for replicates.

For each peach sample, approximately 0.08 to 0.1 g of frozen tissue ground in liquid nitrogen was extracted with 1 mL of cold acetonitrile: isopropanol: water (3:3:2, v/v/v) containing 10 nM of abscisic acid-d6 (Toronto Research Chemicals) and 10 nM salicylic acid-^13^C6 (Santa Cruz Biotechnology) for 3 hours on ice. Extracts were vortexed and spun down for 5 minutes at 13,000 x g. One hundred mL of the extract was then evaporated in a speed evaporator and reconstituted in 100 mL acetonitrile: water (1:9, v/v). Samples for abscisic and salicylic acid measurements were further diluted 10-fold with the extraction solvents, dried, and reconstituted in 100 mL acetonitrile: water (1:9, v/v). Standard calibration curves were constructed using salicylic acid (Cayman Chemicals), indole acetic acid (Cayman Chemicals), and abscisic acid (Milipore-Sigma) over a range of concentrations from 0.98 nM to 1 mM with labeled SA-^13^C6 and ABA-d6 as internal standards. Quantification was done on a Waters Xevo TQ-XS UPLC/MS/MS (Waters Corporation, Milford, MA). Five mL of the sample were separated using an 8-min LC gradient on a Waters Acquity BEH C18 UPLC column (2.1 x 50 mm, 1.7 mm) with mobile phases consisting of water + 0.1% formic acid (solvent A) and methanol (solvent B). The 8 min gradient was: 2% B at 0.00 to 0.10 min, linear gradient to 40% B from 0.10 min to 4.00 min, linear gradient to 70% B from 4.00 to 5.00 min, linear gradient to 99% B from 5.00 to 6.00 min, and hold at 99% B from 6.00 to 7.00 min, then return to 2% B from 7.01 to 8.00 min. The flow rate was 0.4 mL/min and the column temperature was held at 40 °C. The mass spectrometer was operated in both positive and negative electrospray ionization mode and data were collected in multiple reaction monitoring (MRM) channels listed in Supplemental Table S4. Source parameters are as follows: capillary voltage: 1.0 kV; source temperature: 150 °C; desolvation temperature: 400 °C; desolvation gas flow: 800 L/hour; and cone gas flow: 150 L/hour.

Statistical analysis was performed as described for wood composition, with the model, Response= mean + Genotype + Internode + Side + Genotype*Internode + Internode*Side + Internode*Genotype + Internode*Genotype*Side + error, where the response is the level of IAA, SA, or ABA. For comparisons where the ANOVA showed significance for at least one factor, t-tests were used, otherwise a Sidak’s tests were applied.

### RNA Sequencing and read alignment

For RNA sequencing, tissue collection was performed as for the hormone analysis, except each shoot tip was collected separately. All four sample types (IN1 top, IN1 bottom, IN2 top, and IN2 bottom) were sequenced for two shoot tips per tree, for two trees per genotype. Library preparation and sequencing were performed by the Michigan State University Genomics Core. Libraries were created using Illumina Stranded mRNA Prep and indexed with IDT for Illumina RNA UD Indexes. Quality control and quantification for the libraries were performed using a combination of Qubit dsDNA HS and Agilent 4200 TapeStation HS DNA1000 assays. A single pool, with equimolar proportions of each library, was quantified using the Kapa Biosystems Illumina Library Quantification qPCR kit. The pool was loaded onto three lanes of an Illumina HiSeq 4000 single read flow cell and single end 50 bp sequencing was performed using HiSeq 4000 SBS reagents. Base calling was performed using Illumina Real Time Analysis (RTA) v2.7.7 and the output of RTA was demultiplexed and converted to FastQ format with Illumina Bcl2fastq v2.20.0.

FastQ files were processed in CLC Genomics workbench v22. For each library, the data from all three lanes was merged into a single file prior to adapter and quality trimming. Using the trim reads tool set to remove adaptor and following sequence (3’ trim), all files were trimmed for the Illumina Stranded mRNA Prep adaptor (CTGTCTCTTATACACATCT). The mismatch cost =2, gap cost=3, minimal internal match score=10, minimum end match score=4. No homopolymer trimming or filtering based on length. Following adaptor trimming, the trim reads tool was run again to perform quality trimming, with a quality score limit of 0.001 and maximum of 2 ambiguous nucleotides.

Trimmed reads were aligned to the peach genome v. 2.0 (Verde et al., 2017). For RNAseq alignment, CLC Genomics v22.0 the RNA-Seq Analysis (GE) tool was used with parameters as follows: reverse strand specificity, library type= no 3’ bias, mismatch cost=2, insertion cost=3, deletion cost=3, length fraction=0.9, similarity fraction=0.8, with a maximum number of hits=6, expression value=RPKM.

### Transcriptome analysis

For differential expression analysis, the CLC Differential Expression for RNA-Seq tool was used to compare differential expression between upper and lower shoot tip for each genotype by internode combination, controlling for the effect of shoot tip (biological replicate). No filtering was applied prior to FDR calculation.

### Heatmaps and Network Analysis

Genes of interest (GOI) were selected for each internode as those genes that, in either weep or standard, had a Bonferroni of <0.01 and a 2-fold change. This identified 97 GOI for IN1, and 213 GOI for IN2. Heatmaps were created using the ggplot2 package in R. The best arabidopsis homolog of each gene was identified from the Genome Database of the Rosaceae, which uses a blastx algorithm with an expectation value cutoff of 1 e^-6^ (Jung et al., 2019). 81 arabidopsis homologs were identified for the 97 peach IN1 GOI and 164 arabidopsis homologs were identified for the 213 peach IN2 GOI.

To further assess gene interactions and identify functional groups, STRING (v. 11.5) was used to create an interaction network for the arabidopsis homologs with a medium confidence cutoff for interactions (0.4) and a FDR stringency of 5%. The IN1 network had 29 edges, with a p value of 1.79e^-10^. The IN2 network had 449 edges, with a p-value of <1 e^-16^).

To visualize the network, it was imported into Cytoscape (v. 3.9.1). Since multiple peach homologs sometimes mapped to a single arabidopsis gene, the best peach homolog for each arabidopsis gene was manually selected based on the alignment score, and expression values based on the best peach homolog were assigned that arabidopsis homolog. Expression values were mapped to the node color. The protein annotations from STRING were used to manually assign each gene to a functional category, which was mapped to the node outline. Confidence scores for edges ranged from 0.4 to 1 and were mapped to a line width of 0.5 to 8.0.

### Protein Expression Vector Cloning

The *Prunus persica* WEEP coding sequence was PCR amplified using primers JA_211 (5’-CTGTACTTCCAGATGATGATGAGGGAGATGAGCAAAGA-3’)and JA_212 (5’-GTGAGACCACGCAGATGGTTCCAGCTTCAAGGA-3’). The pMal-Strep vector was linearized by PCR using primers JA_205 (5’-TCTGCGTGGTCTCACCCG-3’ and JA_206 (5’-CATCTGGAAGTACAGGTTCTCCCC-3’). The resulting PCR products were assembled using Infusion cloning (Takara) to create the pMal-Strep-WEEP.

### Protein Expression, Purification, and Size Exclusion Chromatography

WEEP protein was expressed with N-terminal 6xHis and maltose binding protein (MBP) tags and a C-terminal StrepTagII. WEEP protein was expressed in *E. coli* NEB Express cells (New England BioLabs). Bacterial cultures were grown at 37 °C to an OD_600_ of ∼0.6, induced with a final concentration of 1 mM isopropyl β-D-1-thiogalactopyranoside (IPTG), then grown overnight at 18 °C. Bacterial cells were pelleted by centrifugation at 5,000 g for 10 minutes, then resuspended in lysis buffer (50 mM tris pH 8.0, 20 mM imidazole, 500 mM NaCl, 10% (v/v) glycerol, 1% Tween-20) supplemented with 1,000 units of lysozyme and 25 units of benzonase nuclease (Millipore Sigma) per ml. Cells were lysed by sonication and cell debris was pelleted by centrifugation at 10,000 g for 45 minutes at 4 °C. The soluble cell lysate was passed through a 0.22 µm filter and then loaded onto to a Ni-NTA chromatography column. The Ni-NTA column was washed with wash buffer (50 mM tris pH 8.0, 20 mM imidazole, 500 mM NaCl, 10% (v/v) glycerol), and bound protein was eluted with elution buffer (50 mM tris pH 8.0, 250 mM imidazole, 500 mM NaCl, 10% (v/v) glycerol). To obtain higher purity full length WEEP protein, the eluted protein was next loaded onto a StrepTactin Sepharose column (IBA Life Sciences). The column was washed with buffer W (100 mM Tris pH 8.0, 150 mM NaCl, 1 mM EDTA) and the bound protein eluted with buffer E (100 mM Tris pH 8.0, 150 mM NaCl, 1 mM EDTA, 2.5 mM desthiobiotin). The eluted protein was concentrated to 1 mg/ml using Amicon Ultra 10 kDa MWCO filters (Millipore Sigma). Size-exclusion chromatography was performed using a Superose 6 10/300 GL column (Cytiva) equilibrated with 1x phosphate buffered saline (PBS) pH 7.4. All protein quantification was done using the Qubit Protein Assay (Invitrogen).

### Root architecture Analysis

Vernalized peach seeds with a radicle emerging collected from field-grown standard and weeping peach trees were planted in root boxes with 1.5” x 2’x 2’ internal dimensions made of opaque white plastic, with a single clear plexiglass side. These were filled with a commercial soil mix without perlite (Sta-Green Tree & Shrub Garden Soil), placed at a 45° angle with the plexiglass side oriented down and covered with black felt, and grown in standard greenhouse conditions. The root system was imaged weekly with a Canon EOS M5 camera from 2 weeks after planting until 10 weeks after planting. Week 4 and week 9 were selected as representative weeks for image analysis for quantitative measurements.

Images were analyzed in RootNav 1.8.1 (Pound et al., 2013; https://sourceforge.net/projects/rootnav/). Some root systems had secondary roots that were co-dominant with the primary root and had tertiary roots branching from them. Because RootNav does not have a setting for tertiary roots, these large secondary roots were identified in RootNav as primary roots, and the tertiary roots off them as secondary. Lateral root number is reported as the total number of secondary and tertiary roots, calculated as (the number of “primary” roots in RootNav - 1) + (the number of “secondary” roots in RootNav). For week 4 data, the entire visible root system was traced, and used to calculate root tip angle, convex hull, and lateral root number. For week 4, primary, secondary, and tertiary roots were used to calculate root tip angle. For the week 9 data, due to the increased complexity of the root system, each image was analyzed twice. First, the entire system was traced and used to calculate convex hull and lateral root number. Second, the main primary root was traced along with a subset of secondary lateral roots which emerged between 80 and 300 mm from the start of the primary root and were visible from emergence to tip. These lateral roots were used to estimate root tip angle and emergence angle. For convex hull and lateral root number, pairwise comparisons between the genotypes were performed using t-tests in Excel, after checking whether the data were normal with equal variances in R (v. 4.1.2 and v.4.2.1). Tests for tip angle, emergence angle, and convex hull were one-tailed, tests for lateral root number were two-tailed. Each test assumed heterogenous or equal variances as appropriate to the data.

### Root Gravitropism Study

Seven standard and seven *weep* peach seedlings approximately six months old were root pruned to about 10 cm of root length, lateral roots were thinned into a single plane, and the shoot was pruned to approximately 30 cm. Seedlings were transplanted into root boxes (constructed as previously described) and allowed to grow for 24 days, until the seedlings had about 15-30 cm of visible new root growth. Boxes were then photographed with a Canon EOS M5 camera and rotated 90°. After rotation they were photographed at 3, 6, and 9 hours, 1-6 days, and again at 16 days.

3-6 roots per plant which were vertically oriented prior to rotation were selected for measurements. Root angle was assessed at 2 days, 4 days, and 6 days. Each root was then assessed for whether they grew and responded to gravity. After excluding roots that failed to grow or were otherwise un-measurable, 50 roots remained for analysis. ImageJ was used to measure the angle between the initial root growth trajectory and the root tip angle trajectory.

Because there were interaction effects between the genotype and the timepoint, the effect of genotype was modeled slicing by timepoint using the lme4 library in R (v. 4.2.1) with blocking by tree (Response= mean + Genotype + Tree:Genotype + error, where Response is the root angle at a particular timepoint, Genotype is the effect of genotype, and Tree is the effect of the individual tree, nested within genotype. The effect of tree was treated as a random variable. Data were assessed for normality and equal variance, then pairwise comparisons between genotypes were performed using emmeans to apply two-tailed t-tests assuming equal variances.

## Acknowledgements

The authors acknowledge Kat Rockwell and Emma Grant for assistance with tree care and imaging and Dr. Miranda Haus for guidance on root analyses. We also would like to acknowledge Mass Spectrometry and Metabolomics Core at Michigan State University, particularly Cassandra Johnny, for troubleshooting, optimizing, and performing the hormone analysis for the peach shoot tips.

## Author Contributions

AK, JLH, JA, LS, FWT, and CAH designed the research. AK, AS, JLH, JA, JA-H, CG, and CAH performed the research and analyzed data. AK and CAH wrote the manuscript.

## Supplemental data

**Supplemental Figure S1.** Neutral sugar content of hand dissected standard (Std) and weeping (weep) tissues from upper and lower regions of each branch. Glucose levels were significantly lower in weeping branch tissues compared to standard branch tissues (p < 0.5). There were no statistically significant differences between tissues or genotypes for all other sugars. Bars represent standard error and n = 4 for each tissue type.

**Supplemental Figure S2.** Longitudinal sections from standard and weeping (weep) peach branches starting at approximately 0.5 cm below the shoot tip. (A) Picture of entire standard and weeping branch sections and zoomed in images of upper (yellow rectangle) and lower (black rectangle) regions of each. Yellow asterisk indicates location of lateral bud in the standard shoot (B) Section of a weeping shoot with a lateral bud. The presence and placement of lateral buds was not predictable or uniform for either genotype, and their presence influenced cell file organization, complicating cell size measurements. Direction towards shoot tip and location of upper (adaxial) side of the branch is indicated. Bar represents 100 µm.

**Supplemental Figure 3.** Hormone concentrations. View of shoot tip dissection for hormone analysis and RNA sequencing. Shoot tips were divided into internodes 1 and 2 (IN1 and IN2), and each internode was bisected into top and bottom (A). The inset shows a cross-section view of the dissection. Each tissue type was tested for auxin (IAA, B), abscisic acid (ABA, C), and salicylic acid (SA, D) Error bars show standard error. All pairwise comparisons done with Sidak’s tests if ANOVA was not significant, or t-tests if ANOVA was significant. Means with the same letter are not significantly different at α=0.05. Bracketed sets indicate the results of paired t-tests between top and bottom within a genotype and internode category, n.s. indicates not significant at the α=0.10 level.

**Supplemental Figure S4.** Principal component analysis for RNAseq data from and lower shoot tissues of internodes (IN) 1 and 2 from both standard (S) and weeping (W) peach branches. Sample naming system indicates genotype (S or W), followed by tree identification number (e.g., 3-2), followed by internode (e.g. IN2 for internode 2), followed by tissue type (i.e., upper or lower).

**Supplemental Figure S5.** Principal component analysis for RNAseq data from upper and lower shoot tissues from internode 1 (Top graph) and internode 2 (Bottom graph) from both standard (S) and weeping (W) peach branches. Sample naming system indicates genotype (S or W), followed by tree identification number (e.g., 3-2), followed by tissue type (i.e., upper or lower).

**Supplemental Figure S6** Uncategorized differentially expressed genes in IN2. Heatmap indicates fold changes between the upper and lower sides of shoots from standard and weeping (weep) trees. Red indicates that the expression is higher on the lower side, blue indicates that the expression is higher on the upper side.

**Supplemental Figure S7.** Differential expression of *SAUR* genes. Heatmap indicates fold changes between the upper and lower sides of shoots from standard and weeping (weep) trees. Red indicates that the expression is higher on the lower side, blue indicates that the expression is higher on the upper side. Grey indicates expression was not detected in that tissue.

**Supplemental Figure S8.** Calibration Curve for SEC flow cytometer and corresponding protein standard information.

**Supplemental Table S1** Differentially expressed genes for IN1, with manual category assignment.

**Supplemental Table S2** Differentially expressed genes for IN2, with manual category assignment.

**Supplemental Table S3** Expression of putative peach *SAUR* genes.

**Supplemental Table S4** Multiple reaction monitoring (MRM) channels of compounds.

