## Supplemental Figures for "Defying Gravity: *WEEP* promotes negative gravitropism in *Prunus persica* (peach) shoots and roots by establishing asymmetric auxin gradients"

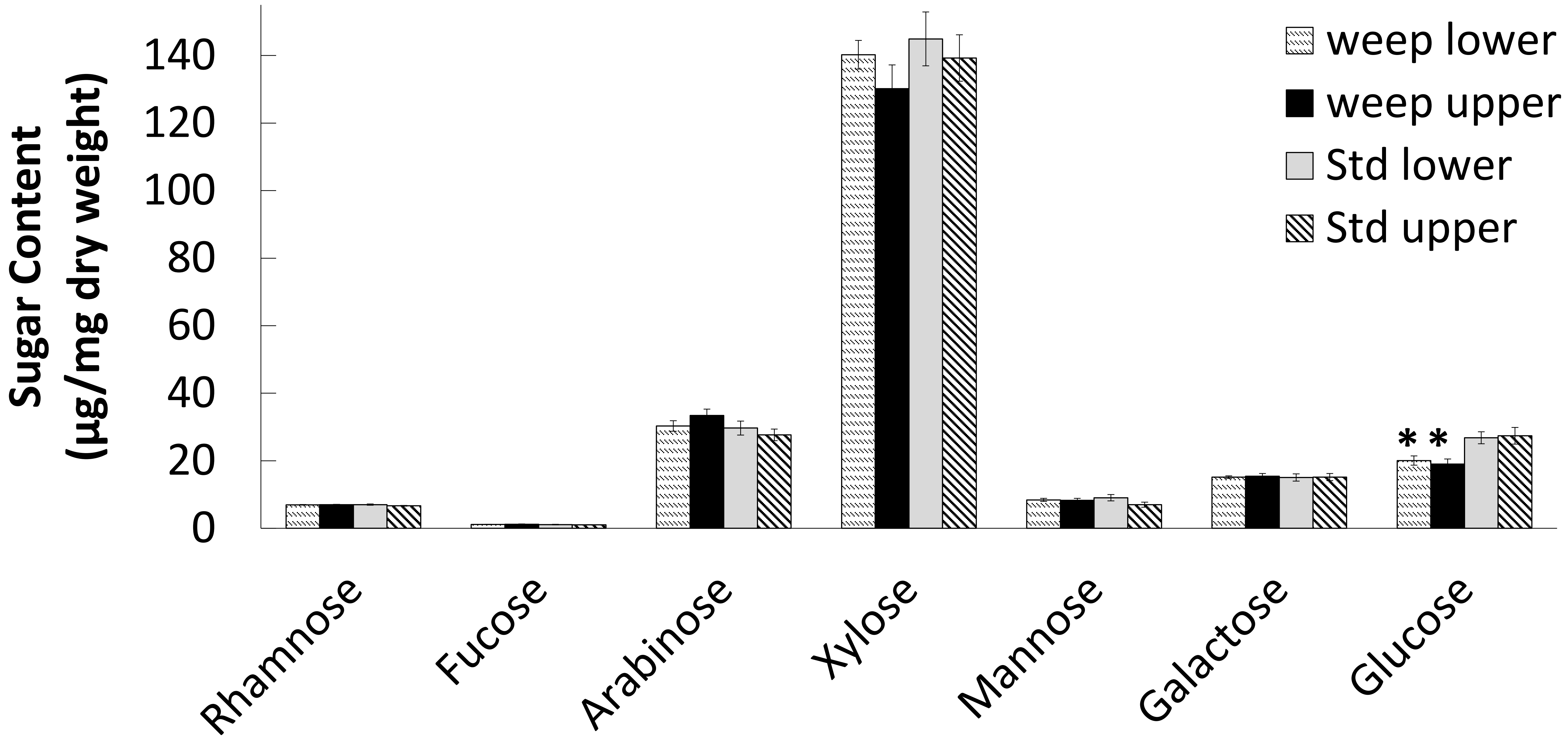


**Supplemental Figure S1**. Neutral sugar content of hand dissected standard (Std) and weeping (weep) tissues from upper and lower regions of each branch. Glucose levels were significantly lower in weeping branch tissues compared to standard branch tissues ($p < 0.5$). There were no statistically significant differences between tissues or genotypes for all other sugars. Bars represent standard error and n = 4 for each tissue type.

.

A

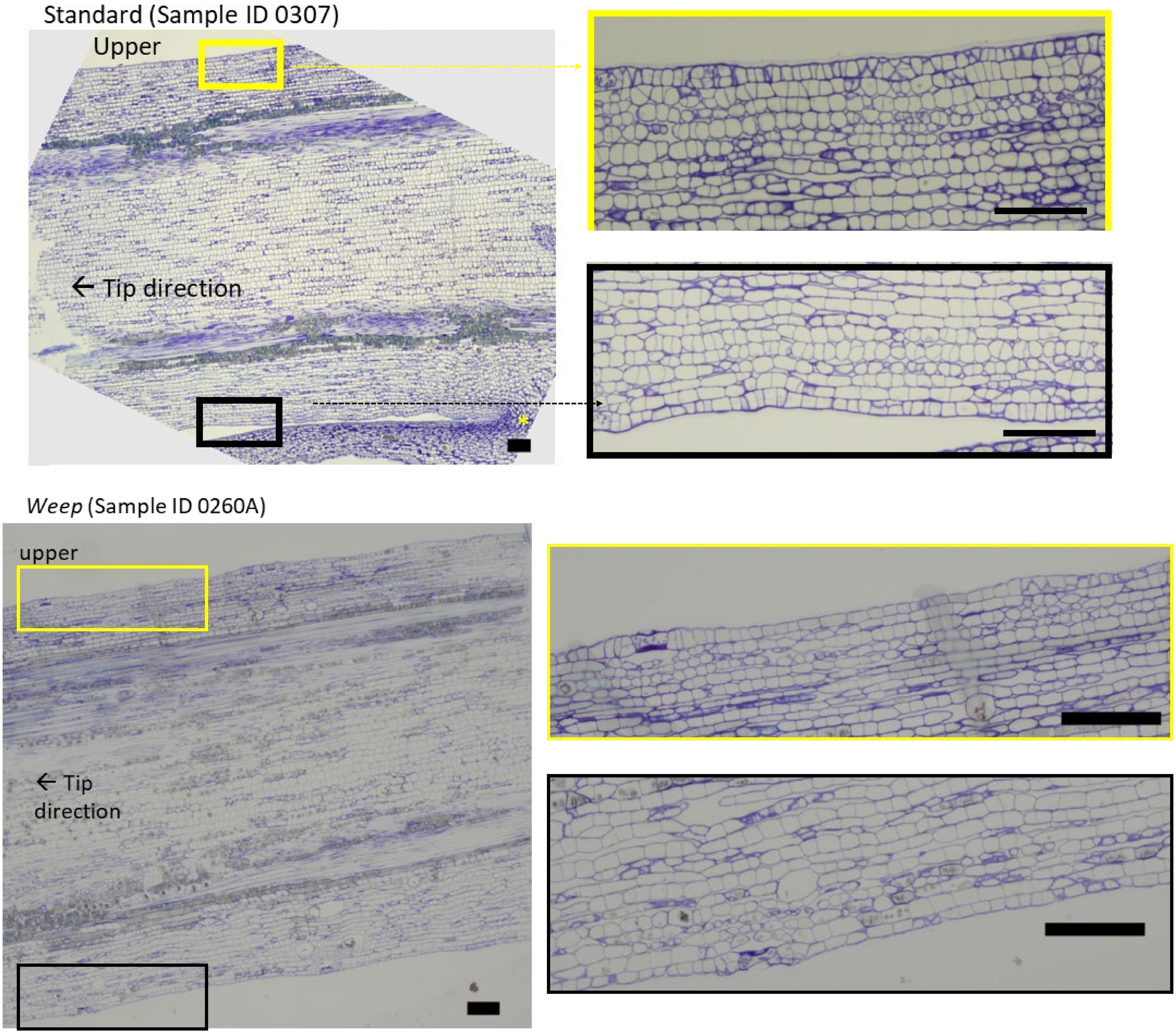


B

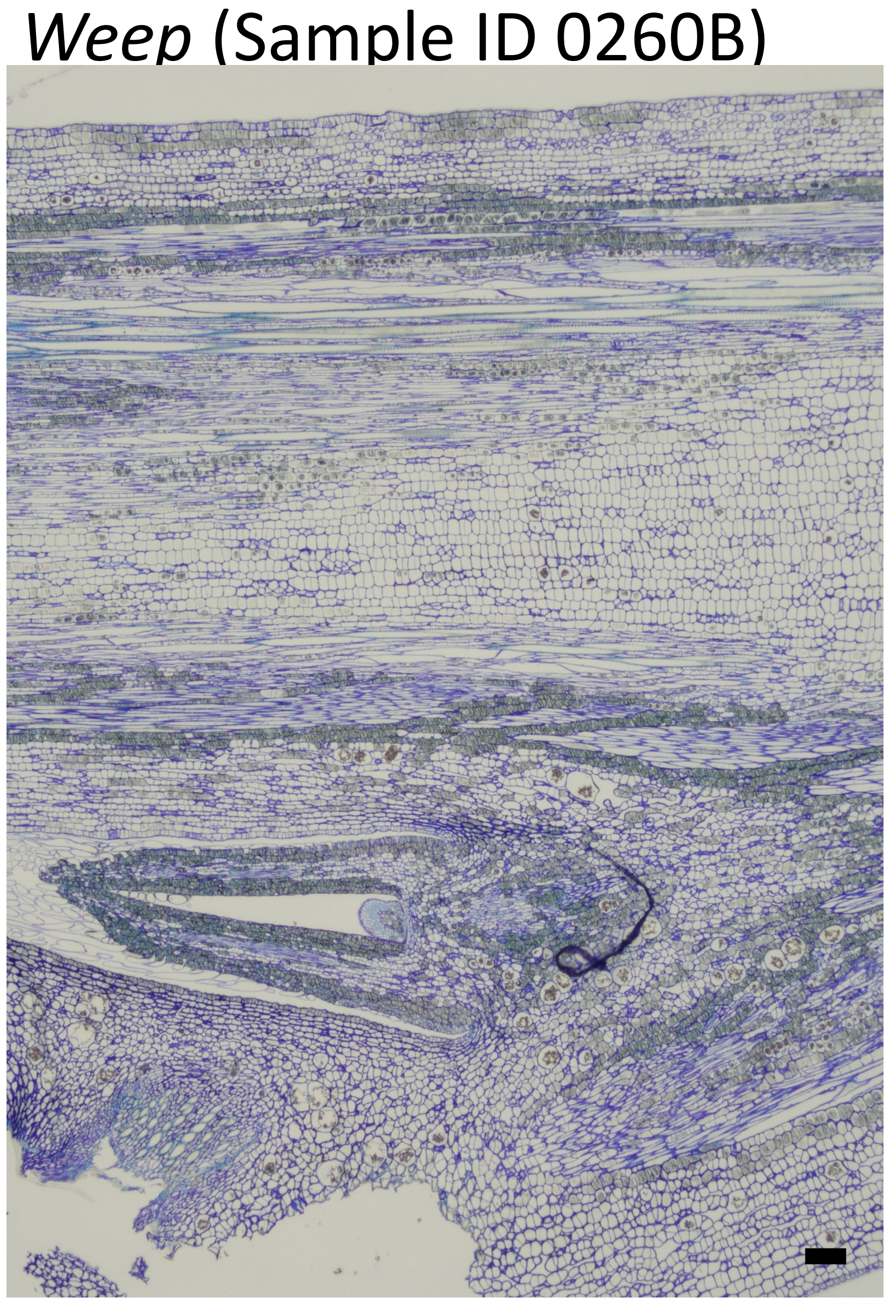


**Supplemental Figure S2**. Longitudinal sections from standard and weeping (weep) peach branches starting at approximately 0.5 cm below the shoot tip. (A) Picture of entire standard and weeping branch sections and zoomed in images of upper (yellow rectangle) and lower (black rectangle) regions of each. Yellow asterisk indicates location of lateral bud in the standard shoot (B) Section of a weeping shoot with a lateral bud. The presence and placement of lateral buds was not predictable or uniform for either genotype, and their presence influenced cell file organization, complicating cell size measurements. Direction towards shoot tip and location of upper (adaxial) side of the branch is indicated. Bar represents 100 µm.

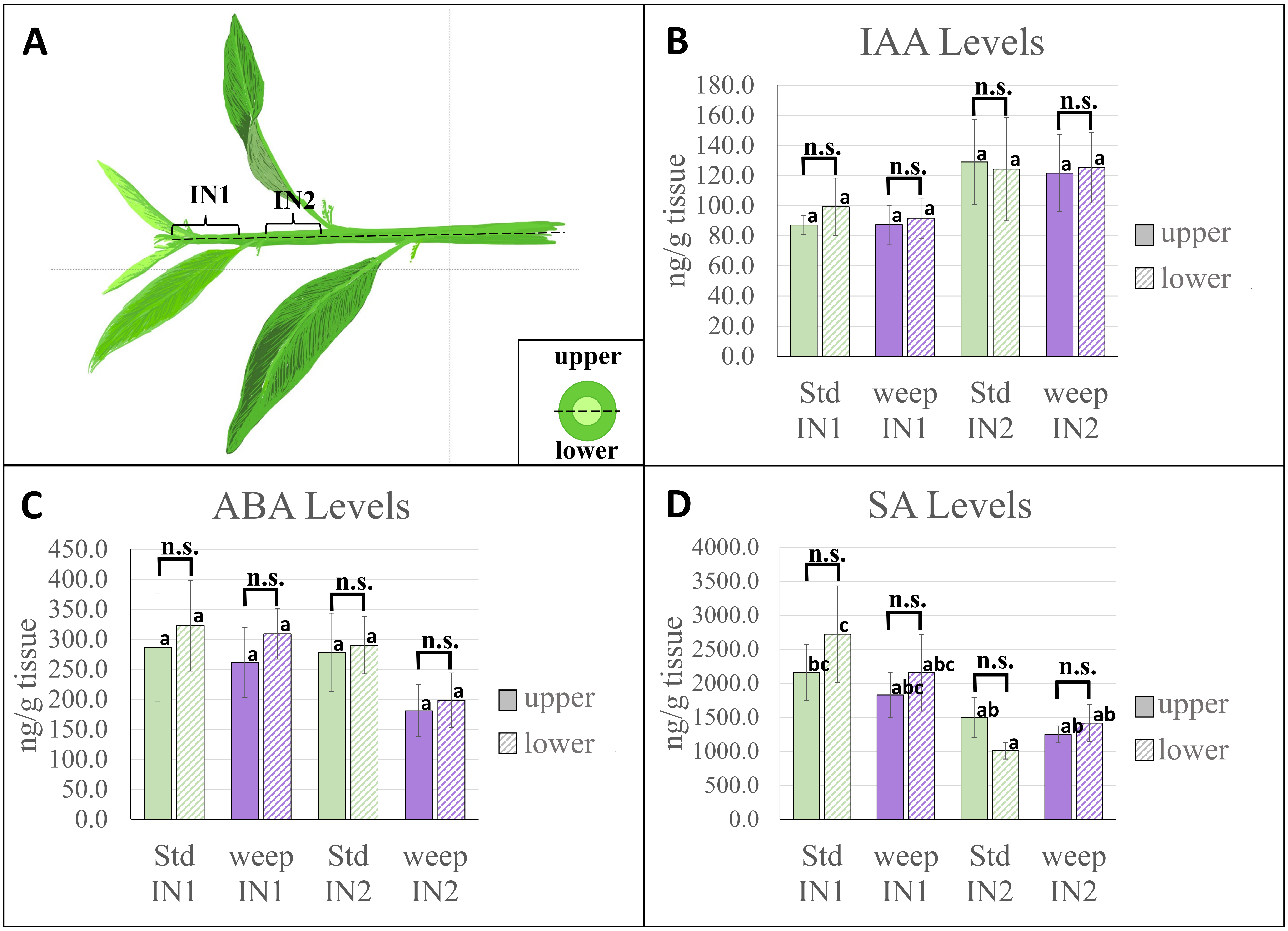


**Supplemental Figure 3.** Hormone concentrations. View of shoot tip dissection for hormone analysis and RNA sequencing. Shoot tips were divided into internodes 1 and 2 (IN1 and IN2), and each internode was bisected into top and bottom (A). The inset shows a cross-section view of the dissection. Each tissue type was tested for auxin (IAA, B), abscisic acid (ABA, C), and salicylic acid (SA, D) Error bars show standard error. All pairwise comparisons done with Sidak's tests if ANOVA was not significant, or t-tests if ANOVA was significant. Means with the same letter are not significantly different at α=0.05. Bracketed sets indicate the results of paired t-tests between top and bottom within a genotype and internode category, n.s. indicates not significant at the α=0.10 level.

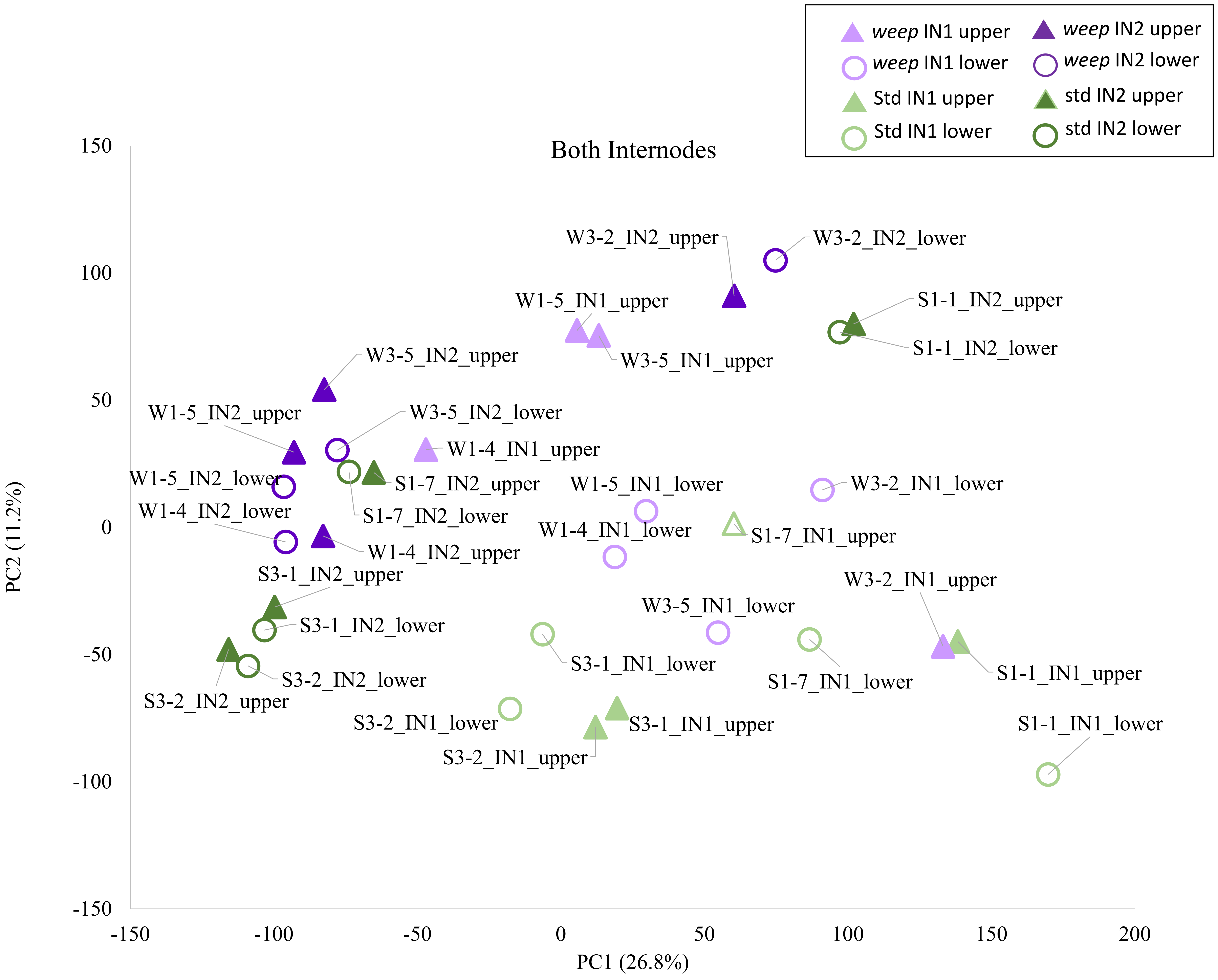


**Supplemental Figure S4.** Principal component analysis for RNAseq data from and lower shoot tissues of internodes (IN) 1 and 2 from both standard (S) and weeping (W) peach branches. Sample naming system indicates genotype (S or W), followed by tree identification number (e.g., 3-2), followed by internode (e.g. IN2 for internode 2), followed by tissue type (i.e., upper or lower).

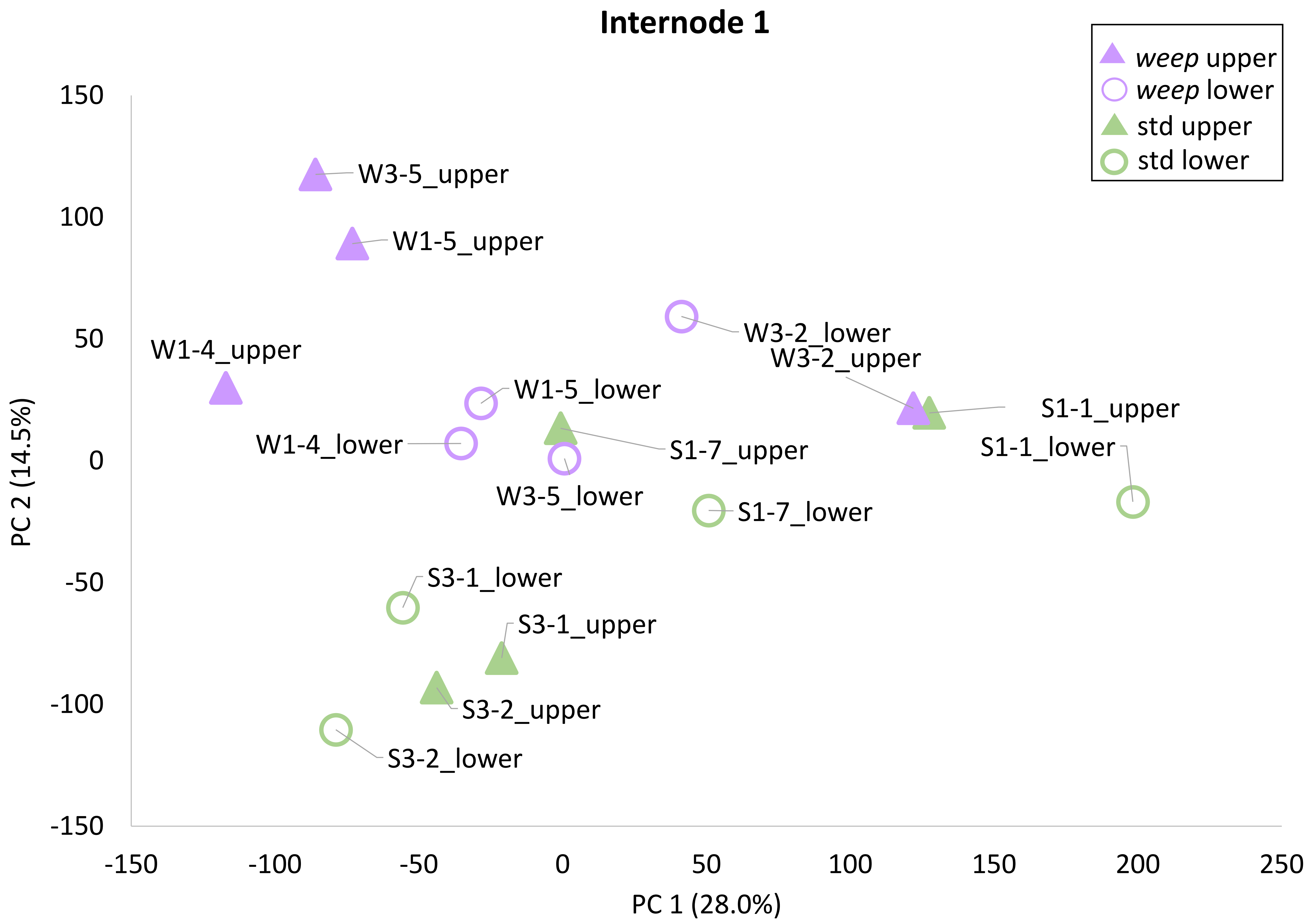


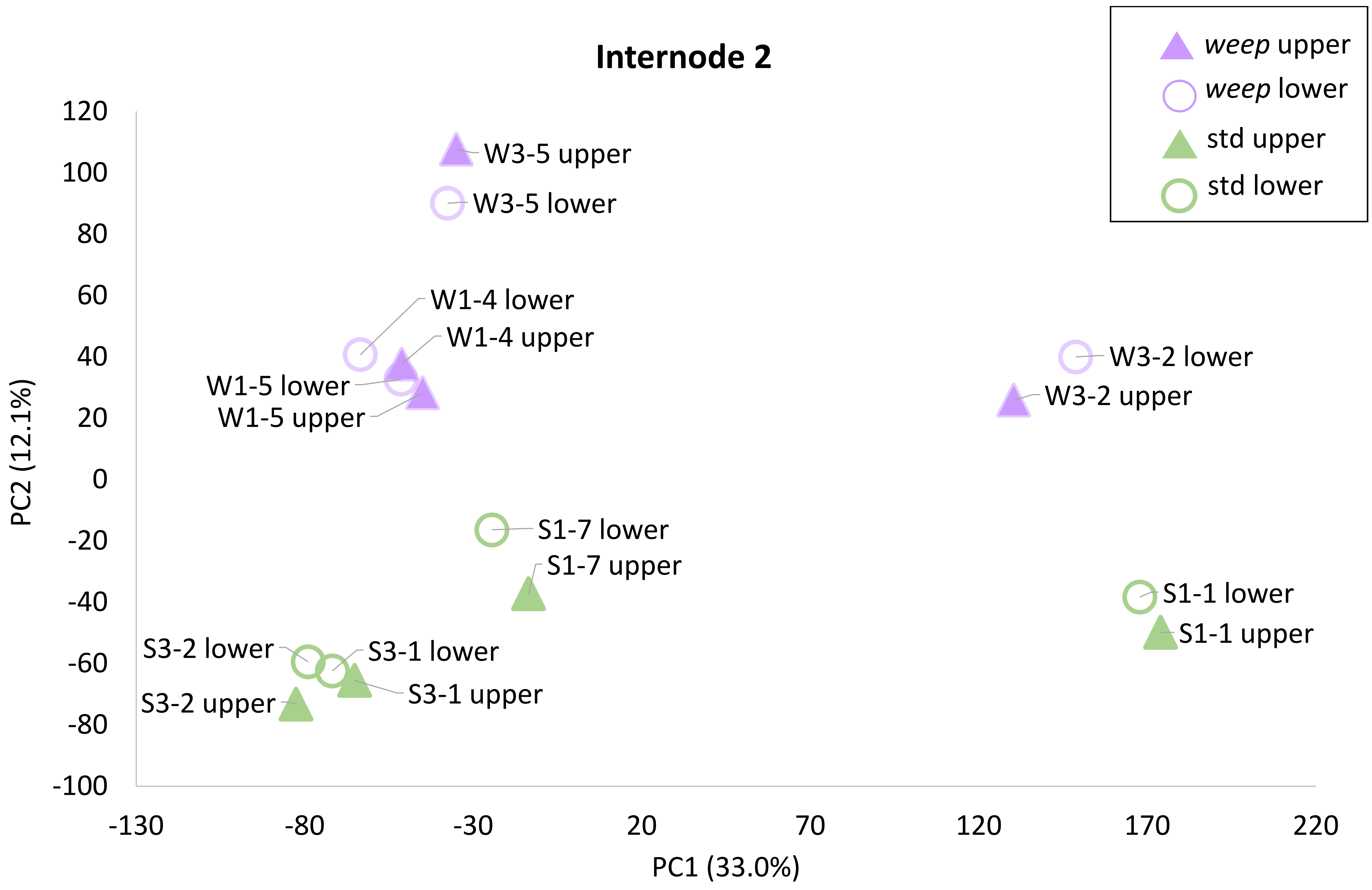


**Supplemental Figure S5.** Principal component analysis for RNAseq data from upper and lower shoot tissues from internode 1 (Top graph) and internode 2 (Bottom graph) from both standard (S) and weeping (W) peach branches. Sample naming system indicates genotype (S or W), followed by tree identification number (e.g., 3-2), followed by tissue type (i.e., upper or lower).

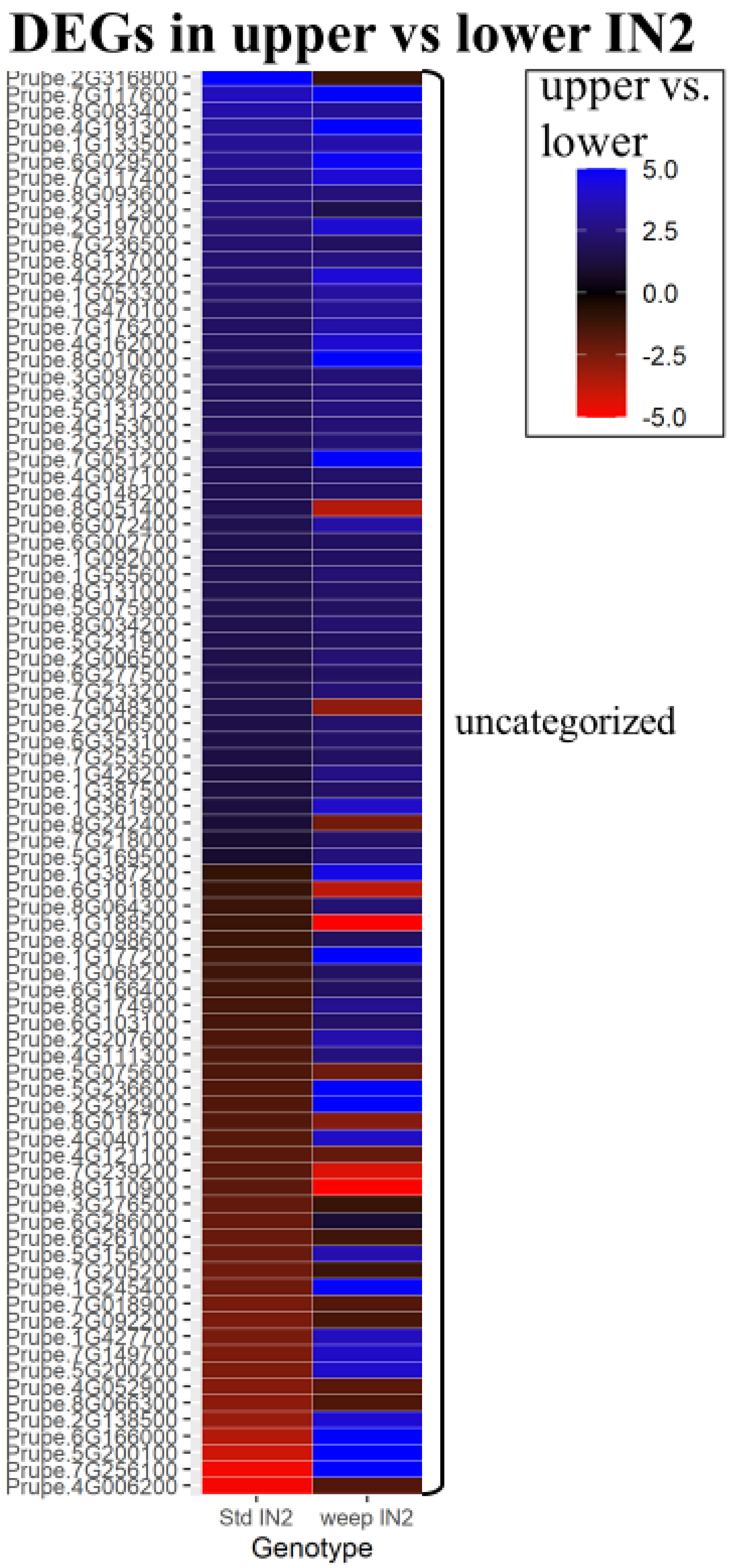


**Supplemental Figure S6** Uncategorized differentially expressed genes in IN2. Heatmap indicates fold changes between the upper and lower sides of shoots from standard and weeping (weep) trees. Red indicates that the expression is higher on the lower side, blue indicates that the expression is higher on the upper side.

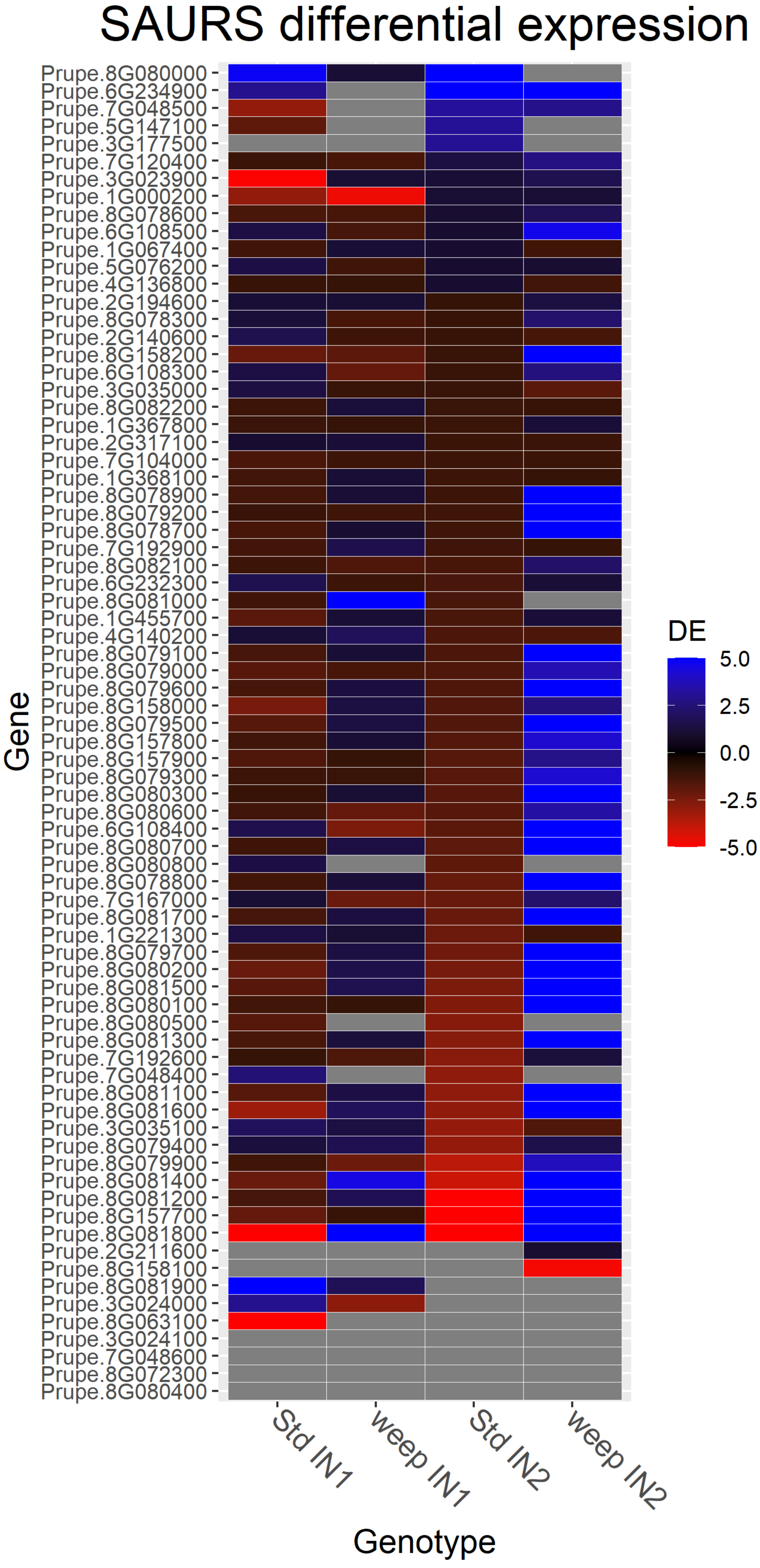


**Supplemental Figure S7.** Differential expression of *SAUR* genes**.** Heatmap indicates fold changes between the upper and lower sides of shoots from standard and weeping (weep) trees. Red indicates that the expression is higher on the lower side, blue indicates that the expression is higher on the upper side. Grey indicates expression was not detected in that tissue.

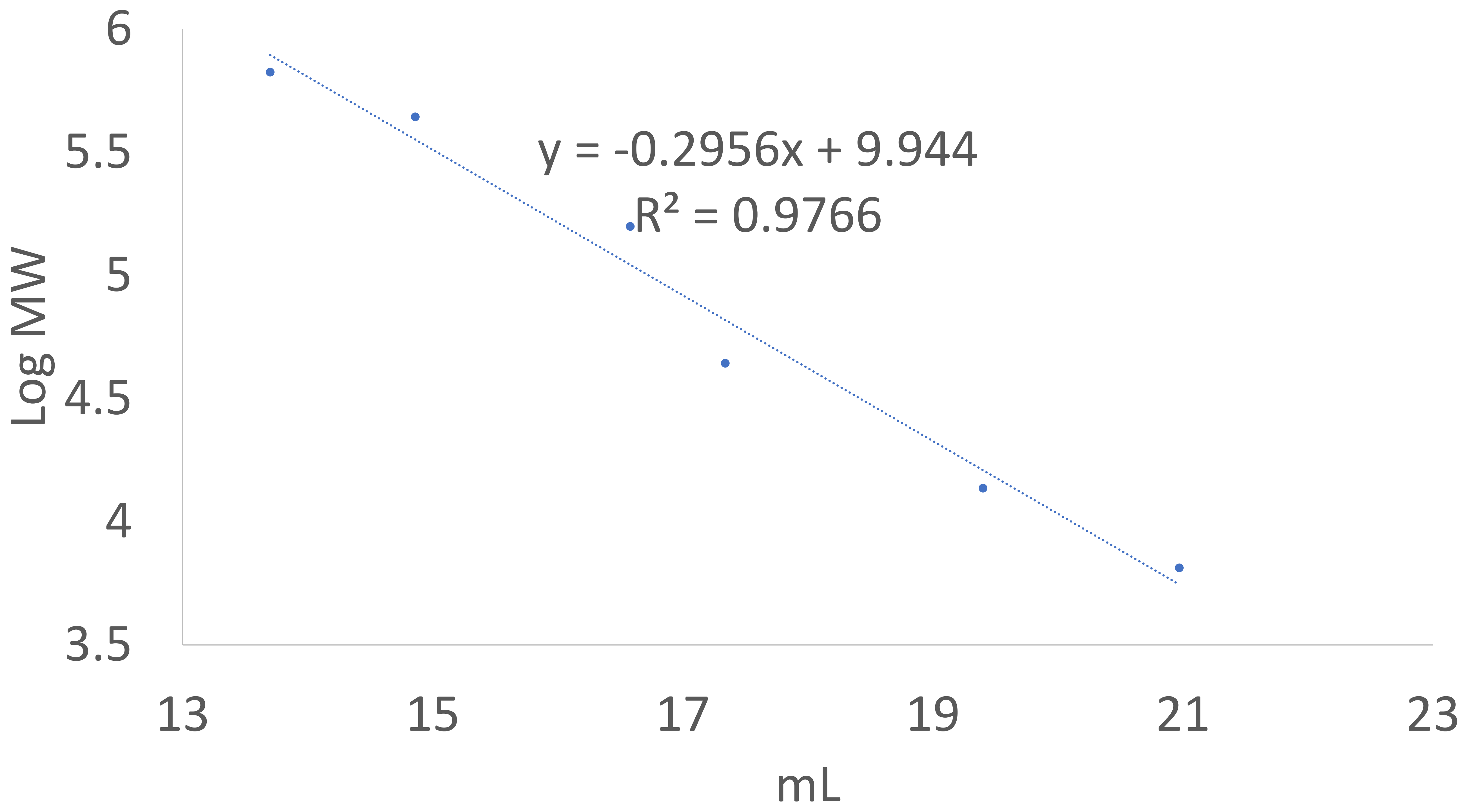


| Standard | Ve | LogMW | MW |
|---|---|---|---|
| Aprotinin | 20.97 | 3.81291336 | 6500 |
| Ribonuclease A | 19.4 | 4.13672057 | 13700 |
| Ovalbumin | 17.34 | 4.64345268 | 44000 |
| Aldolase | 16.58 | 5.19865709 | 158000 |
| Ferritin | 14.86 | 5.64345268 | 440000 |
| Thyroglobulin | 13.7 | 5.82542612 | 669000 |

**Supplemental Figure S8.** Calibration Curve for SEC flow cytometer and corresponding protein standard information.
